# Single-cell atlas of human liver development reveals pathways directing hepatic cell fates

**DOI:** 10.1101/2022.03.08.482299

**Authors:** Brandon T. Wesley, Alexander D. B. Ross, Daniele Muraro, Zhichao Miao, Sarah Saxton, Rute A. Tomaz, Carola M. Morell, Katherine Ridley, Ekaterini D. Zacharis, Sandra Petrus-Reurer, Judith Kraiczy, Krishnaa T. Mahbubani, Stephanie Brown, Jose Garcia-Bernardo, Clara Alsinet, Daniel Gaffney, Olivia C. Tysoe, Rachel A. Botting, Emily Stephenson, Dorin-Mirel Popescu, Sonya MacParland, Gary Bader, Ian D. McGilvray, Daniel Ortmann, Fotios Sampaziotis, Kourosh Saeb-Parsy, Muzlifah Haniffa, Kelly R. Stevens, Matthias Zilbauer, Sarah A. Teichmann, Ludovic Vallier

## Abstract

The liver has been studied extensively due to the broad number of diseases affecting its vital functions. However, therapeutic advances, especially in regenerative medicine, are currently hampered by the lack of knowledge concerning human hepatic cell development. Here, we addressed this limitation by describing the developmental trajectories of different cell types comprising the human fetal liver at single-cell resolution. These transcriptomic analyses revealed that sequential cell-to-cell interactions direct functional maturation of hepatocytes, with non-parenchymal cells playing critical, supportive roles during organogenesis. We utilised this information to derive bipotential hepatoblast organoids and then exploited this novel model system to validate the importance of key signalling pathways and developmental cues. Furthermore, these insights into hepatic maturation enabled the identification of stage-specific transcription factors to improve the functionality of hepatocyte-like cells generated from human pluripotent stem cells. Thus, our study establishes a new platform to investigate the basic mechanisms of human liver development and to produce cell types for clinical applications.

---

The liver fulfils a broad spectrum of functions including blood detoxification, metabolite storage, lipid/glucose metabolism and secretion of serum proteins. These critical tasks are mainly performed by the hepatocytes which are supported by a diversity of cell types. Kupffer cells are tissue-specific resident macrophages responsible for liver homeostasis and immunity^1^. Hepatic stellate cells sequester vitamin A in healthy organs while promoting fibrosis through collagen secretion during disease^2^. Cholangiocytes form the epithelial lining of the biliary tree, which transports bile into the intestine^3^, and play a role in liver repair during chronic injury^4, 5^. Finally, sinusoidal endothelial cells provide a permeable interface with circulating blood and promote regeneration after liver damage^6^. Importantly, adult liver cells have been broadly characterised using diverse methods, including detailed single-cell transcriptomic analyses^7–10^. However, the study of these cell types during fetal life remains limited, especially in humans due to there being only a handful of studies that are descriptive in their nature^11^. This knowledge gap presents a major challenge in the advancement of new therapies including regenerative medicine applications. Here, we address this limitation by performing single-cell RNA sequencing (scRNA-seq) analyses on human fetal and adult livers (Fig. 1a). This single-cell map not only uncovered the developmental trajectories of the different cell types comprising the liver, but also the cell-cell interactions controlling organogenesis. We also took advantage of this information to isolate human hepatoblasts, which represent the early progenitors of the liver proper, and demonstrated that they can be propagated as organoids to model developmental processes. Finally, we utilised this map to assess the differentiation path of human pluripotent stem cells (hPSCs) into hepatocyte-like cells (HLCs) and uncovered transcription factors capable of improving the resemblance of HLCs to adult hepatocytes. Together our results present novel insights into liver development which allow the establishment of an *in vitro* platform for modelling human liver development while providing the knowledge necessary to improve the production of hepatocytes *in vitro*^12^.

**Fig. 1.**
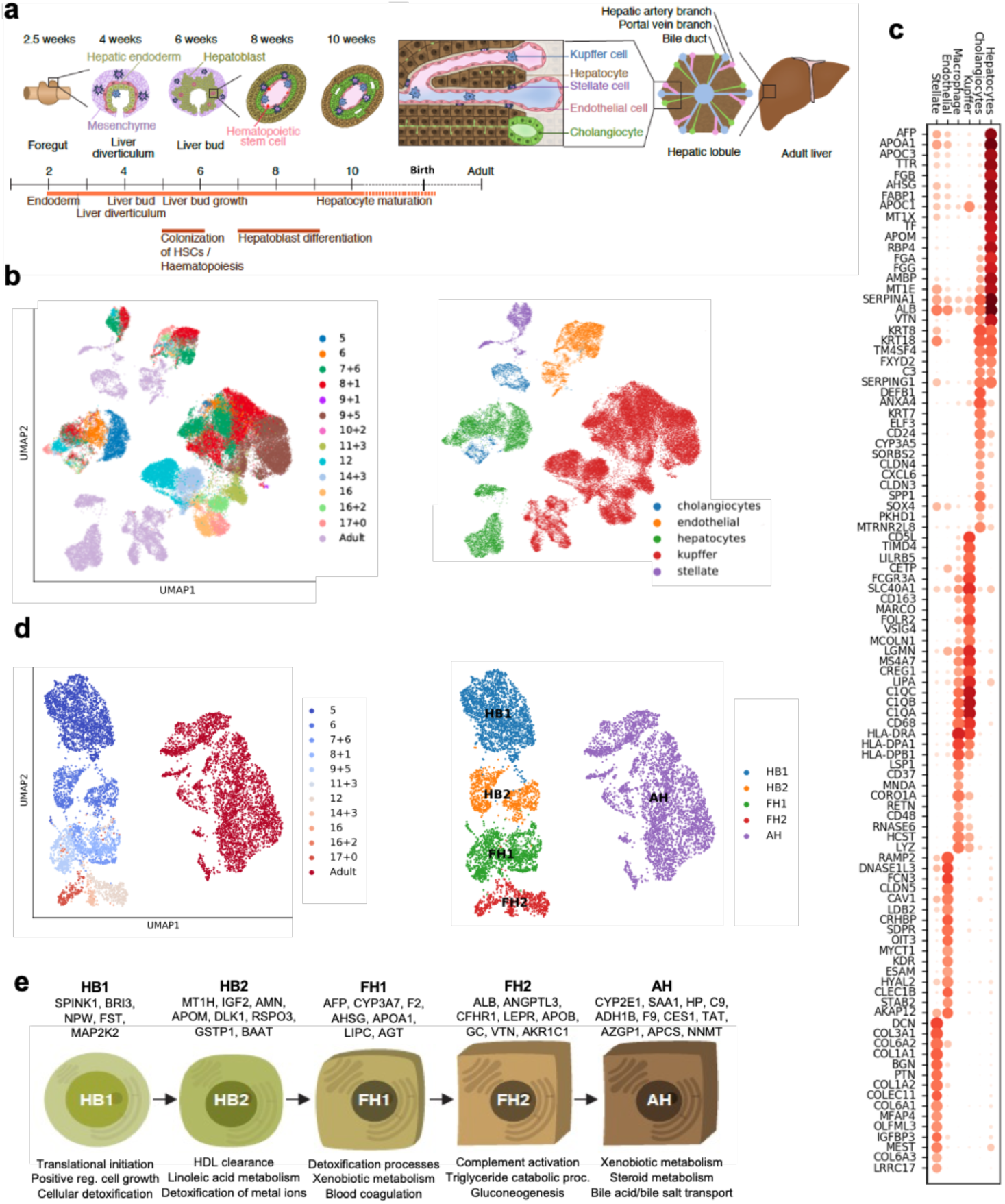
Single-cell transcriptomic map of human liver development. **a,** Schematic representation of human liver development. **b,** UMAP visualization of fetal and adult human hepatic cells from 10x Genomics sequencing, annotation indicates post-conceptional weeks (PCW) + days (left panel) and the different lineages (right panel). **c,** Mean scaled, log-normalized gene expression values of selected differentially expressed genes (DEGs) for each hepatic cell lineages. Gene-expression frequency (fraction of cells within each cell type expressing the gene) is indicated by dot size and level of expression by colour intensity. **d,** UMAP visualization of hepatocyte developmental trajectory (left panel) and annotation of developmental stages based on Louvain analysis (right panel): HB1, hepatoblast stage 1; HB2, hepatoblast stage 2; FH1, fetal hepatocyte stage 1; FH2, fetal hepatocyte stage 2; AH, adult hepatocyte. **e,** Specific genes induced at each stage of hepatocyte differentiation and corresponding gene ontology (GO).

## Single-cell transcriptomic map of the developing human liver

To characterise the cellular landscape of the developing liver (Fig. 1a), human single-cells were collected using methods tailored to each stage of development or extracted from existing data sets (Supplementary Table 1). Droplet-based single-cell RNA-sequencing was performed to profile a total of 237,978 hepatic cells, of which 87% passed quality control (see Methods). UMAP dimensionality reduction and sub-clustering of these cell’s transcriptome showed that our approach captured the main cell types comprising the liver (Fig. 1b,c and Supplementary Table 1). Of note, cholangiocytes were the least represented cell type in our collection confirming the difficulty of isolating these cells from liver tissue^8^. In addition, cholangiocytes were only identified at 7 post-conception weeks (PCW) reinforcing previous studies indicating that these cells differentiate from hepatoblasts after 7-8 PCW^13, 14^. Concerning endothelial cells, the first cells were captured from 5 PCW at the time when liver vasculature is known to be established^15, 16^. Tissue-resident Kupffer cells could be distinguished from monocyte-derived macrophages by the expression of MARCO, CD163, FCGR3A and CD5L^8^ and the absence of LSP1 and CD48 (Fig. 1c). Finally, hepatic stellate cells were captured from 5 PCW, supporting studies in mice suggesting that these cells could be derived from the septum transversum at 3-5 PCW^2, 17, 18^. Of note, all data generated by this study can be easily viewed here: https://collections.cellatlas.io/liver-development. Importantly, transcriptomic observations were validated by immunostaining on primary human fetal liver (Extended Data Fig. 1a). Collectively, these results show that our single-cell atlas captured the major cell types of the liver and their dynamic diversity during development.

## Single cell RNA-Sequencing reveals the developmental trajectory of liver cells

Utilizing this dataset, we examined the developmental trajectory of each cell type, starting with hepatocytes. Principal component analysis (PCA), Louvain clustering and diffusion pseudotime (dpt) analyses defined 5 hepatocyte developmental stages (Fig. 1d and Extended Data Fig. 1b): hepatoblast stages 1 and 2 (HB1 at 5 PCW and HB2 at 6 PCW), fetal hepatocyte stages 1 and 2 (FH1 at 7-11 PCW and FH2 at 12-17 PCW) and adult hepatocytes (AH). Each stage displayed distinct transcriptional changes indicative of unique cell states (Extended Data Fig. 1c-e and Supplementary Table 3). Accordingly, this analysis identified markers specific to each stage including SPINK1 for hepatoblasts, GSTA1 for fetal hepatocytes, and hepatoglobin (HP) for adult hepatocytes. We also observed that each stage of hepatocyte development was marked by the induction of genes associated with specific liver function (Fig. 1e). Thus, hepatocytes follow a progressive functional maturation during organogenesis corresponding to the acquisition of hepatic activity during fetal life.

We then performed similar analyses on cholangiocytes, stellate cells, endothelial cells, and Kupffer cells (Extended Data Fig. 2 a-h and Supplementary Tables 4-7). Briefly, only cholangiocytes seemed to gradually differentiate from the HB2 stage, whereas PCA analyses did not reveal major differences among sequential timepoints for most non-parenchymal hepatic cell types. More precisely, Louvain clustering and diffusion pseudotime allowed the distinction of an embryonic stage at 5-6 PCW, intermediate fetal stage between 7-17 PCW, and an adult state (Extended Data Fig. 2a-h). This suggested that these cell types may not undergo a significant functional maturation during fetal life after their initial embryonic specification. Interestingly, these three stages correspond to major modifications in the liver environment: liver bud formation, colonisation by the haematopoietic system at 7 PCW and shift from fetal to adult cells^16^. Thus, these data suggest that the developmental trajectory of hepatic cells is influenced by major developmental events while only hepatocytes seem to undergo a progressive functional maturation.

We then decided to further demonstrate the interest of our single cell map to define the embryonic origin of specific cell types. We decided to focus on hepatic stellate cells since previous studies have reached divergent conclusions^2^. Louvain clustering on early stellate cells revealed a population of cells expressing mesenchymal markers, one population with an endothelial-bias, and a third population combining markers for both lineages (endothelial: CDH5, LYVE1, KDR and STAB2 and stellate cells: PDGFRB, VIM, DES and COL1A1; Fig. 2a-c). Diffusion pseudotime confirmed that fetal endothelial and stellate cells could originate from this stellate-endothelial progenitor population, termed SEpro (Fig. 2 c-e and Supplementary Table 8), while gene expression analyses reveal that these cells expressed genes associated with proliferation and DNA replication characteristic of a stem cell or progenitor state. Immunohistochemistry validations on primary tissue revealed cells expressing both PDGFRB and CDH5 located within the vasculature of the 6 PCW liver (Fig. 2b), thereby confirming the existence of this progenitor *in vivo*. Of note, previous studies in model organisms have suggested the existence of such progenitors without functional demonstration^19, 20^. To further address this limitation, we decided to validate the existence of such progenitors during differentiation of hPSCs in vitro. We first performed scRNA-Seq analyses on hPSCs differentiating into endothelial^21^ and hepatic stellate cells^22^. UMAP, PCA and diffusion pseudotime comparison of these differentiations reveal an overlapping stage sharing the expression of markers specific for both lineages (Extended Data Fig. 3a-d). To confirm that this stage could include a common progenitor, hPSC were differentiated into endothelial cells for 3.5 days and then grown in culture conditions inductive for hepatic stellate cells. The resulting cells were able to transition away from the endothelial pathway characterised by the expression of CDH5, KDR and VWF while acquiring the stellate cells markers PDGFRA, COL1A1, ACTA2, and NCAM (Extended Data Fig. 3e). Taken together, these results illustrate how single cells observations can be combined with in vitro differentiation to further understand the developmental process leading to stellate cells production.

**Fig. 2.**
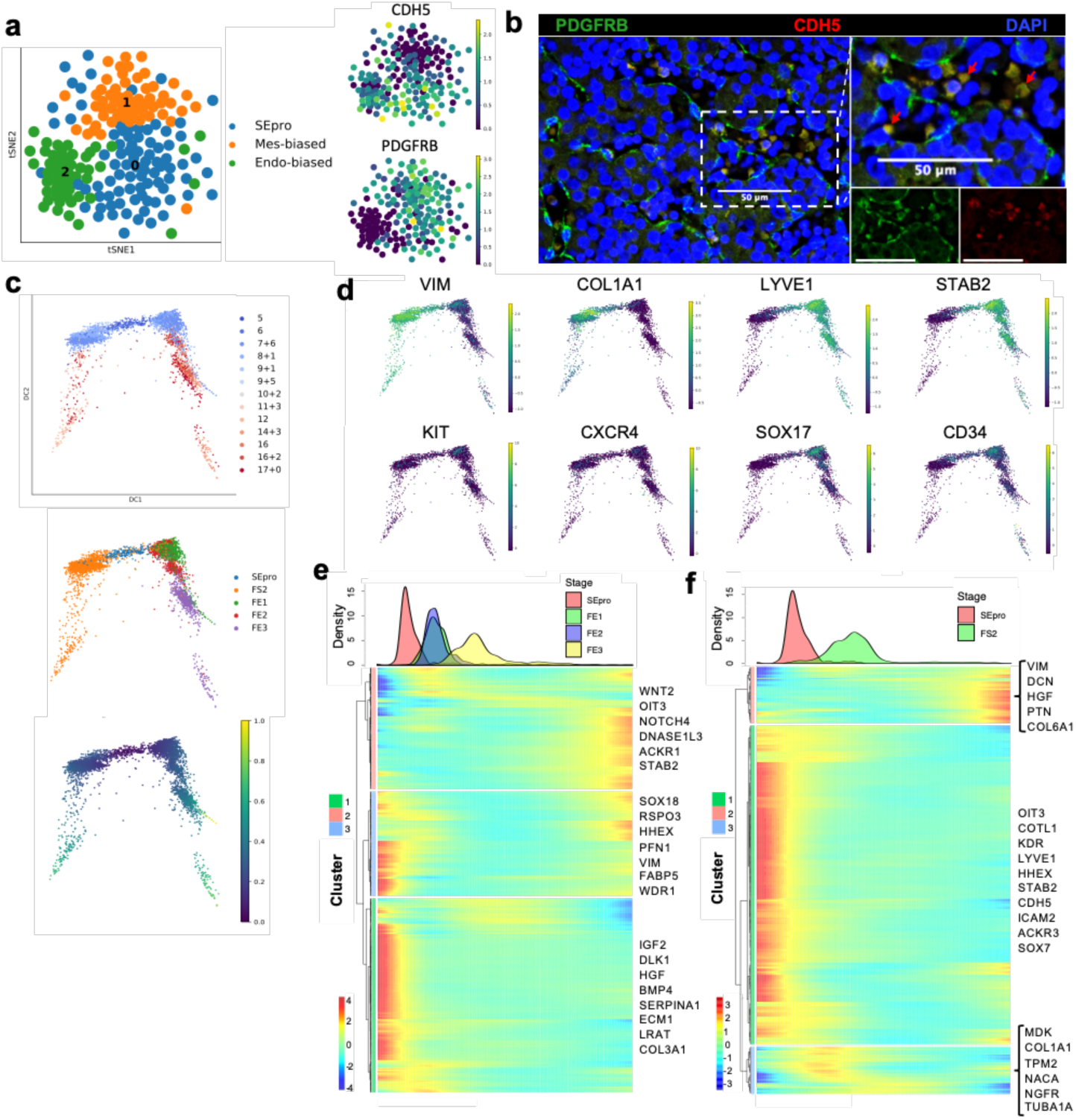
Identification of a hepatic stellate and endothelial cell progenitor in the early fetal liver. **a,** tSNE visualization based on Louvain clustering of 6 PCW human fetal liver cells identifying stellate-endothelial progenitors or “SEpro” (left panel). Gene expression tSNE plots show the co-expression of specific markers for both lineage by SEpros (right panel). **b,** Immunofluorescence staining (IF) of 6 PCW human liver identifying the SEpro population based on co-expression of stellate (PDGFRB) and endothelial (CDH5) markers. Scale bars = 50 um. **c,** Diffusion pseudotime analyses of stellate and endothelial cells developmental trajectories showing that each lineage originated from SEpro. **d,** Diffusion pseudotime analyses of specific markers for each lineage (top row stellate cells, bottom row endothelial cells). **e,** Heatmap of time-related genes during fetal endothelial cell development and **f,** fetal stellate cell development starting with SEpro and progressing toward 17 PCW.

## Human hepatoblasts can be grown *in vitro* while maintaining their differentiation capacity

We then decided to use our single-cell analyses to isolate and grow *in vitro* hepatoblasts, as they represent the natural stem cell of the liver during development. Our single cell analysis showed that hepatoblast HB1 and HB2 display characteristics of early these liver stem cells. Indeed, these cells expressed WNT target genes associated with adult stem cells such as LGR5, high levels of cell cycle regulators suggesting self-renewal capacity, and markers specific for both biliary and hepatocytic lineages indicative of bi-potential capacity of differentiation (Extended Data Fig. 1e). Based on these observations and previous reports showing a crucial role for WNT^23^, we hypothesised that this signalling could support the growth of hepatoblasts *in vitro*. To explore this possibility, 6 PCW livers were dissociated into single cells which were sorted based on EPCAM expression and grown in 3D culture conditions supplemented with WNT (Fig. 3a). The isolated cells formed branching organoids which could be expanded for more than 20 passages (Extended Data Fig. 4a-b). These hepatoblast organoids (HBO) homogeneously expressed hepatoblast markers (Fig. 3b and Extended Data Fig. 4c-d) and single cell RNA-seq analyses demonstrated that they closely resembled their *in vivo* counterparts, especially the HB2 stage (Fig. 3c, Extended Data Fig. 4e- f, and Supplementary Table 9). Of note, neither HBO nor HB1/2 cells express NCAM thereby excluding the presence of hepatic stem cells in our analyses. To confirm HBO bipotentiality, we transplanted tdTomato-HBO into Fah-/Rag2-/Il2rg- (FRG) mice^24^ using an approach developed for primary hepatocytes^25^. After 27 days, implants were recovered, and red fluorescent cells could be observed in all the grafts (Extended Data Fig. 4g) indicating that HBO had engrafted efficiently. H&E staining of explanted tissue sections revealed the presence of numerous nodules resembling densely packed hepatocytes, as well as biliary epithelial-like cells assembled into structures resembling bile ducts (Extended Data Fig. 4h). Engrafted organoids stained positive for both KRT18 and AFP at the time of implant, but AFP was markedly decreased by day 27 (Fig. 3d) suggesting differentiation into hepatocytes *in vivo*. Accordingly, numerous cells in hepatic nodules stained positively for ARG1, A1AT, and ALB while significant levels of human albumin were identified in mouse serum suggesting functional activity of implanted organoids (Fig. 3e). Finally, some KRT18+ nodules were found to contain cells expressing KRT19 (Fig. 3f), either as a mixed population or as a pure KRT19+ population. Together, our results demonstrate that HB2 hepatoblasts can be grown *in vitro* while maintaining their capacity to differentiate into hepatocytes and cholangiocytes.

**Fig. 3.**
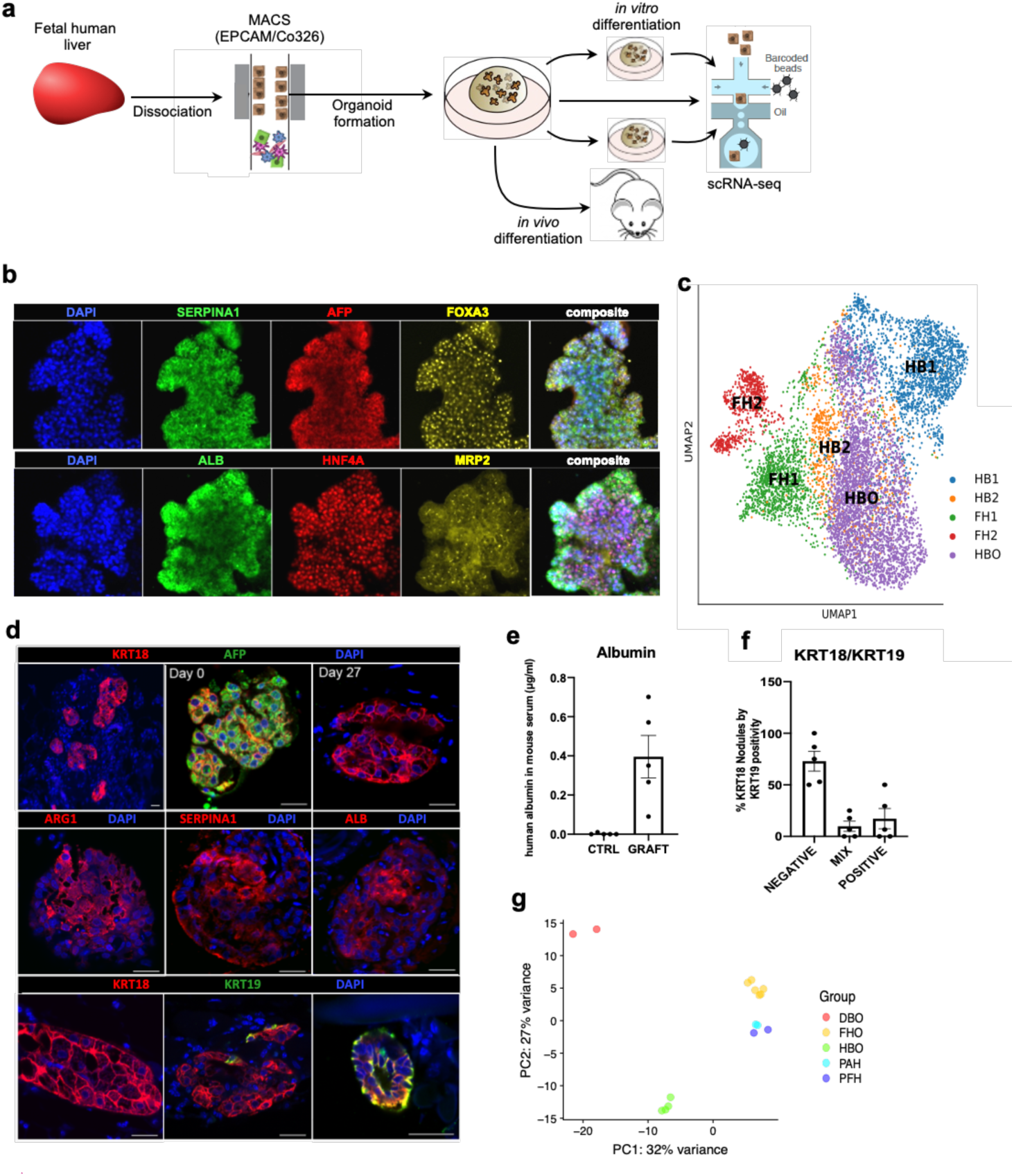
Modelling early hepatic development *in vitro* using novel hepatoblast organoids. **a,** Schematic representation of hepatoblast organoid (HBO) derivation and subsequent analyses. **b,** Immunostaining showing expression of hepatoblast markers in HBO grown in vitro. **c,** UMAP visualisation of fetal hepatoblast/hepatocyte differentiation stages along with HBO confirming that HBO share the transcriptional profile of HB2 stage of hepatocyte development. **d,** IF showing decrease of the foetal hepatocyte marker (AFP) in organoid after 27 days of engraftment while hepatocyte markers (KRT18, ALB, ARG1, and SERPINA1) were maintained suggesting differentiation into mature hepatocytes. IF for biliary markers identified KRT19-positive cells in a subset of nodules, which organised into bile duct-like structures. Unless otherwise stated, pictures show grafts 27 days transplantation. Scale bars = 20 um. **e,** ELISA analyses showing secretion human ALB in the serum of HBO recipient mice 27 days after engraftment. **f,** Quantification (percentage) of KRT19-positive cells within KRT18-positive nodules. **g,** Principal component analysis (PCA) showing the divergence in gene expression profile between hepatoblast organoids (HBO, n=4 lines derived from 4 different fetal livers), differentiated biliary organoids (DBO, n =2), fetal hepatocyte organoids (FHO; n = 6), primary adult hepatocytes (PAH, n=2), and primary fetal liver (PFH, n=2).

Of note, two types of human liver organoid systems have been described previously by Huch et al. (2015)^26^ and Hu et al. (2018)^24^. The former is composed of intrahepatic cholangiocytes which can differentiate towards hepatocyte-like cells^26, 27^ (differentiated biliary organoids or DBO), whilst the latter derives organoids from hepatocytes^24^. Therefore, we characterised both systems against HBO (Fig. 3g). Organoids derived from intrahepatic cholangiocytes expressed markers such as KRT19, but did not express hepatocyte markers in either the undifferentiated nor differentiated state (Extended Data Fig. 4i-j). Transcriptomic comparison also demonstrated the transcriptional divergence between HB2/HBO, DBO and hepatocyte organoids (Fig. 3g and Supplementary Table 9). Analysis of the genes driving this separation revealed hepatocyte and biliary markers, with HBO having intermediate levels of both these sets of markers (Extended Data Fig. 4i-l). Taken together, these data demonstrate that our single-cell analyses have identified a unique self-renewing population of hepatoblasts that can be propagated long-term *in vitro*.

## Dynamic intercellular interactions of the developing liver

To further understand the mechanisms directing liver organogenesis, we captured the interactions between the hepatoblasts/hepatocytes and other cell types using the CellPhone database (CellPhoneDB)^28^ (Fig. 4a, Extended Data Fig. 5 and Supplementary Tables 10-11). This approach revealed that most interactions begin in hepatoblasts, stabilise in fetal hepatocytes and finally disappear in adult cells (Extended Data Fig. 5a). Furthermore, a diversity of unknown interactions was captured between stellate cells, Kupffer cells and endothelial cells indicating potential roles in extracellular matrix organisation, haematopoietic development and innate immunity (Extended Data Fig. 5a). Thus, our analysis could reveal the source of signalling pathways controlling liver development (Supplementary Tables 10-11). Of note, some of these interactions were validated using RNA-Scope on primary tissues (Extended Fig. 5b,c). As an example, NOTCH4/DLL4 was expressed by endothelial cells and could interact with DLK1/NOTCH2 on hepatoblasts/hepatocytes (Fig. 4b and Extended Data Fig. 5c). These bidirectional interactions suggest that hepatoblasts could be involved in the vascularisation of the liver and thus could direct the construction of their own niche (Supplementary Tables 10-11). In return, endothelial cells could control hepatoblast differentiation into cholangiocytes, a process known to require NOTCH signalling^29^. Similarly, RSPO3-LGR4/5 interactions were detected between hepatoblasts and stellate cells at 5-6 PCW (Fig. 4a-b). Thus, stellate cells could support hepatoblast self-renewal by boosting WNT signalling. To confirm this hypothesis, HBO were co-cultured *in vitro* with the hepatic stellate cell line LX2^30^ in the presence or absence of WNT signalling. Expression of proliferation (MKI67), hepatoblast (TBX3) and WNT target (LGR5) genes were upregulated in HBO co-cultured with LX2 cells (Extended Data Fig. 5d,e). However, this effect was variable, suggesting that additional cell types could be required to support HBO self-renewal. This result could also be explained by the activated nature of LX2 cells and by the challenge to replicate short distance cellular interactions in our 3D culture system. Nonetheless, these results validate the relevance of the cellular interactions revealed by our single cell analyses.

**Fig. 4.**
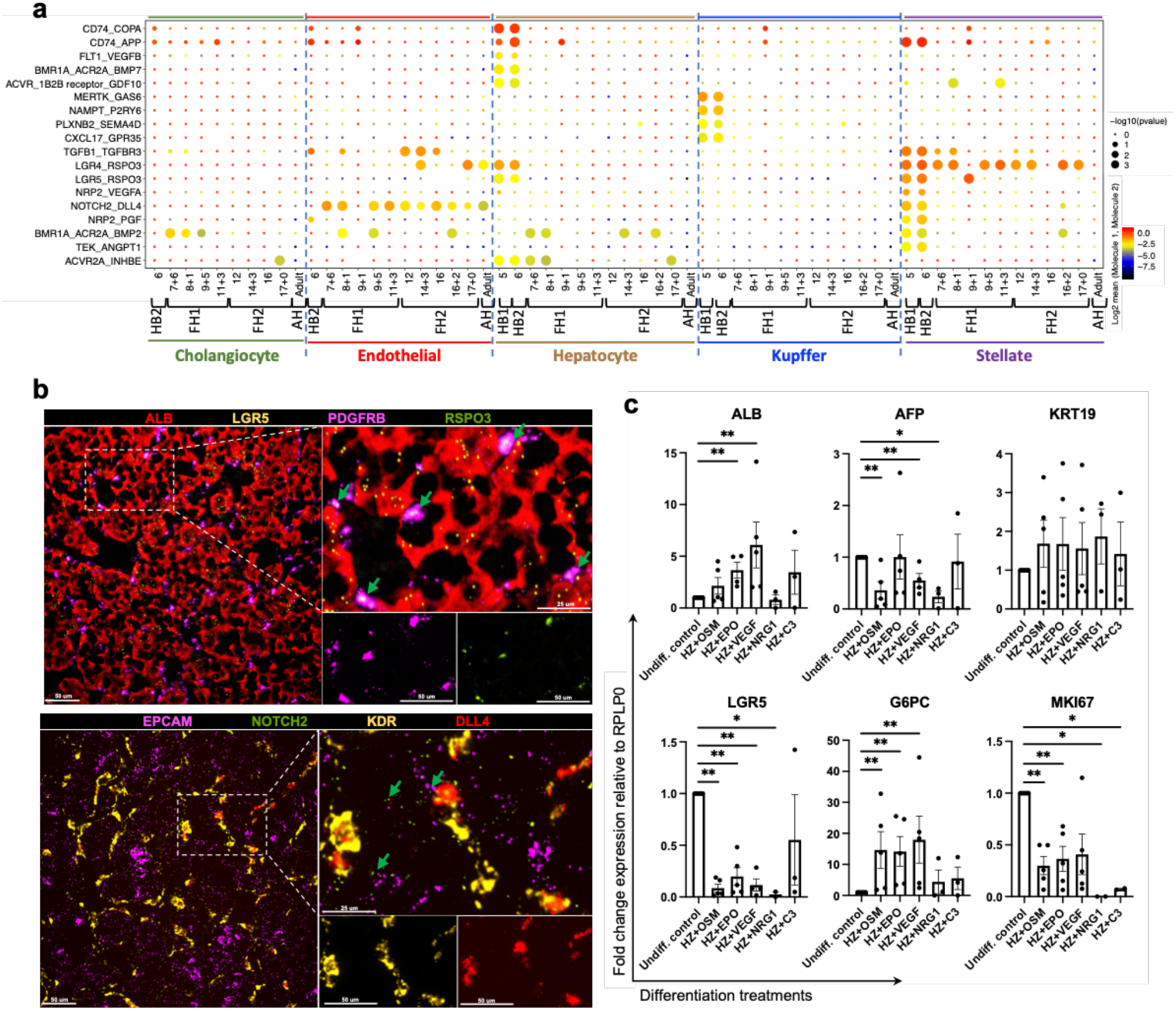
Cell-to-cell interaction networks during human liver development. **a,** CellphoneDB analysis of hepatocytes receptors-ligand interactions with other hepatic cells across all developmental timepoints. Y-axis shows ligand-receptor/receptor-ligand interactions, with the hepatocyte protein listed first; x-axis shows developmental timeline of each cell type; colour shows log2 mean of interacting molecules; size of dot shows -log10(P) significance. **b,** RNAscope validating ligand-receptor interactions establishing the niche of hepatoblast in 6 PWC liver. RSPO3 is expressed hepatic stellate cells while receptor LGR5 is expressed by hepatoblasts (top panel). DLL4 is expressed in from endothelial cells while NOTCH2 receptor is expressed on hepatoblasts (bottom panel). **c,** Quantitative PCR showing the expression of hepatocyte maturation genes following treatment of key signalling molecules discovered using the single-cell liver development atlas. Undiff control: HBO grown in culture conditions maintaining their selfrenewal. HZ : HepatoZYME basal medium. *P<0.05, **P<0.01, ***P<0.001; Mann-Whitney two-tailed, unpaired t-tests.

We then decided to test if these analyses could also be used to identify new pathways controlling hepatocyte and cholangicoyte differentiation. For the former, we selected 5 growth factors suggested by cell phone DB analyses (EGF, VEGFA, C3 NRG1 and EPO, Oncostatin-M or OSM) (Fig. 4a, Extended Data Fig. 5 and Supplementary Tables 10-11). The effect of these factors was then analysed on HBO grown in the absence of WNT to allow their differentiation. Several factors (EPO, OSM and VEGFA) were associated with a decrease in hepatoblast markers (AFP, LGR5 and MKI67) and increase in hepatocyte markers (albumin and G6PC) (Fig. 4c and extended Fig. 6a) suggesting a differentiation toward foetal hepatocytes. Other factors either blocked the expression of hepatocytes markers (NRG1) or have limited effect (C3). To further reinforce these observations, we decided to characterise the effect of OSM by performing scRNA-Seq on HBO induced to differentiate into hepatocytes. These analyses confirmed the transcriptional shift of HBO toward hepatocyte after treatment with OSM (Extended Data Fig. 6b,c and Supplementary Table 12). Furthermore, this differentiation was associated with gain of hepatocyte functions including cytochrome P450 activity (Extended Data Fig. 6d) and lipid accumulation (Extended Data Fig. 6e). Together, these data confirm that HBO can differentiate into hepatocytes in the absence of WNT upon stimulation of specific factors such as OSM.

We next focused on cholangiocytes differentiation. CellphoneDB analyses reinforced previous reports^4, 5^ suggesting an important function for TGFb in this process^31^ (Fig. 4a, Supplementary Table 10). To test this hypothesis, we supplemented HBO media with TGFb for seven days and observed a marked switch toward cholangiocyte identity illustrated by the induction of biliary markers KRT19 and loss of hepatoblast markers (Extended Data Fig. 6d,f-h and Supplementary Table 13). These observations were confirmed by scRNASeq analyses showing that the transcriptome of HBO grown in the presence of TGFb resemble that of cholangiocyte organoids (Extended Data Fig. 6d,i,k) Taken together, these data confirm the interest of our single cell analyses to identify cell-cell interactions directing liver development and also the interest of HBO for validating the function of the signalling pathways involved.

## Liver development atlas informs the maturation of hiPSC-derivatives

To further exploit our single-cell map and the data generated above, we decided to address a key challenge associated with human pluripotent stem cells (hPSCs)^32^ differentiation. It is well established that most protocols currently available to differentiate hPSCs result in cells with fetal characteristics^33^ rather than fully functional, adult-like cells. Accordingly, single-cell analyses and detailed characterisations have demonstrated the fetal identity of hPSC-derived hepatocytes^34, 35^, however, the mechanisms blocking progress toward an adult phenotype remain unclear. To address this question, we performed scRNA-Seq on hPSCs differentiating into hepatocyte-like cells (Fig. 5a). PCA analyses confirmed the progressive process driving the acquisition of a hepatocytic identity (Fig. 5b,c and Supplementary Table 14). This differentiation trajectory was then compared to the developmental trajectory of primary hepatoblasts/hepatocytes by diffusion pseudotime alignment (Fig. 5d), UMAP, PAGA analyses and PCA (Extended Data Fig. 7a-c). These comparisons showed that HLCs at day 14 of differentiation aligned to the second hepatoblast stage, after which their differentiation follows an *in vitro* specific process. Differential gene expression analyses yielded a list of genes related to xenobiotic metabolism, bile acid transport, and lipid metabolism pathways, which suggests that the divergence between HLCs and primary cells prevents the acquisition of fully adult function (Supplementary Table 15). Importantly, a similar divergence from primary development was observed in other cell types generated from hPSCs including cholangiocytes^36^, endothelial cells^21, 37^, stellate cells^22^ and macrophages^38^ (Extended Data Fig. 7d-g; Supplementary Tables 16-19). Thus, hPSC differentiation could systematically deviate from a natural developmental path after embryonic stages preventing the production of adult cells.

**Fig. 5.**
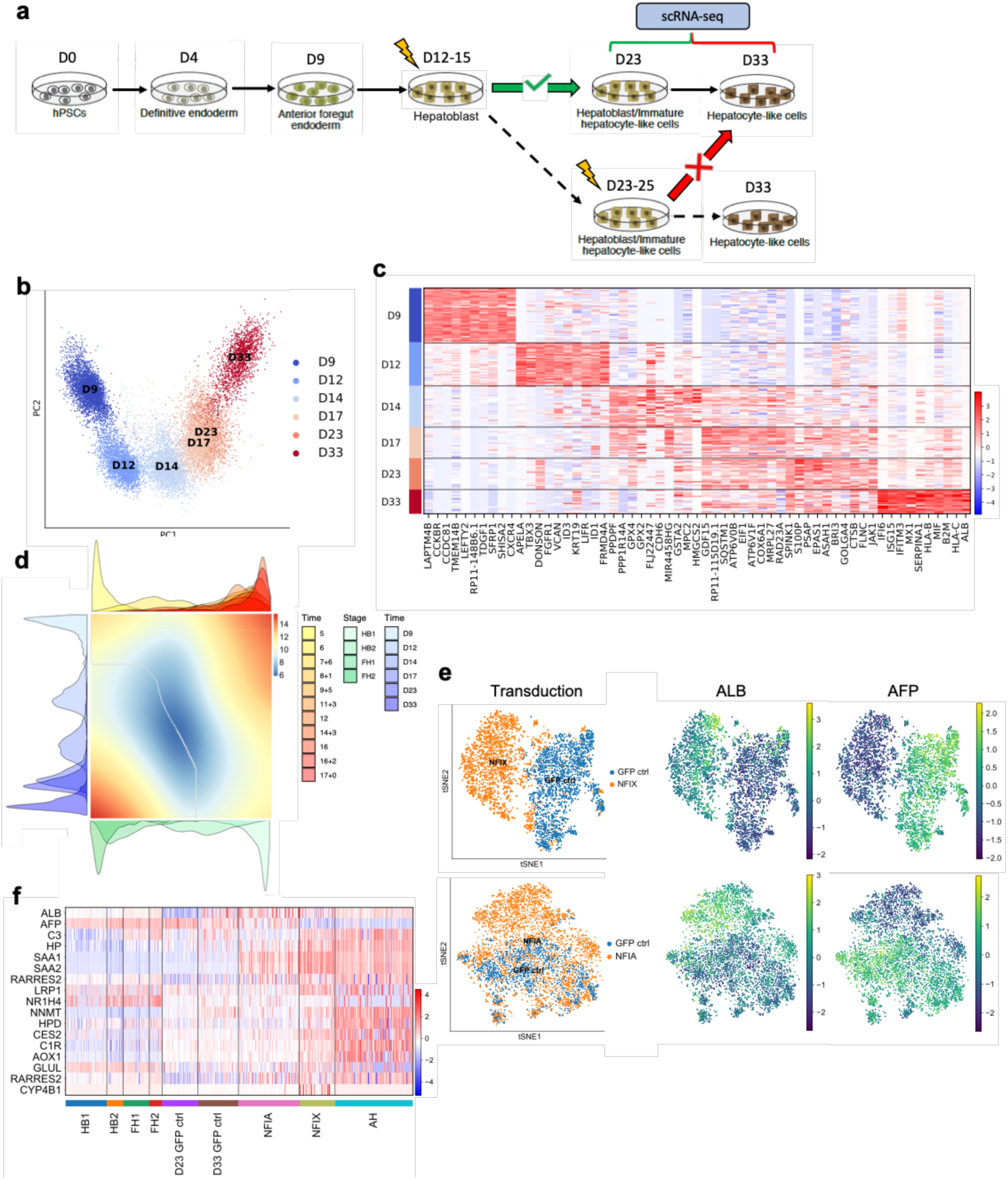
Temporal overexpression of key transcription factors in hPSC-derived hepatocytes increases their similarity to adult primary hepatocytes. a, Schematic representation of experimental processes to validate the function transcription factors in hepatocytes differentiation. b, PCA analyses showing the step-by-step differentiation of hPSCs into hepatocytes. D, day of differentiation. c, Heatmap of top 10 differentially expressed genes (DEG) specific between each stage of differentiation (Wilcoxon-Rank-Sum test, z-score>10). d, Alignment of primary hepatocyte developmental trajectory to hiPSC differentiation using the CellAlign package; red plot shows regions of misalignment/dissimilarity; blue plot shows regions of close alignment/similarity. e, UMAP visualization of HLCs transduced with transcription factors NFIX, NFIA and GFP (control) showing that TFs can increase ALB expression while decreasing the expression of the foetal marker AFP. f, Heatmap showing the acquisition of functional hepatocytes markers in transduced Hepatocytes derived from hPSCs.

Comparison of *in vivo* to *in vitro* hepatocyte differentiation also revealed transcription factors expressed in fetal hepatocytes (FH1) that were missing in hPSC-derived cells (Extended Data Fig. 8a-c; Supplementary Table 15). To validate their functional relevance, these factors were overexpressed in hPSCs differentiated into HLCs for 15 days. The phenotype of the resulting cells was assayed by scRNA-seq 8 days after transduction (Extended Data Fig. 8d; Supplementary Table 20). Of particular interest, NFIX or NFIA expression changed the transcriptional profile of HLCs (Fig. 5e). This shift was characterised by a decrease in fetal markers and the induction of markers indicative of the adult state (ALB, HP, C3; Fig. 5e-f). Furthermore, pathway enrichment analyses showed an increase in specific functions associated with adult hepatocyte identity, including metabolic and complement-related pathways (Extended Data Fig. 8e). Thus, NFIX or NFIA overexpression during differentiation of hIPSCs appear to increase the expression of specific markers likely downstream of these transcritption factors. These results were further validated by inducing the expression of NFIX, NFIA, and CAR during differentiation of hPSCs toward HLCs using the Opti-OX system^39, 40^. Stable induction of NFIX and to a lesser extent NFIA upregulated an array of functional markers including ALB, SAA1, SAA2, LRP1, CES2, AOX1 while downregulating AFP (Extended Data Fig. 9). Taken together, these results show that mis-expression of key developmental regulators during HLC differentiation could explain their limited capacity to become adult hepatocytes, and that expression of these factors at an early step of differentiation could augment their similarity to adult cells.

## Discussion

Our study provides a detailed map of human liver development and shows that the developmental trajectory of liver cells is influenced by changes in liver environment. Colonisation by the haematopoietic system^41^ appears to initiate the functional maturation of hepatocytes, which progressively acquire intrinsic hepatic functions. Of note, the activation of the liver after birth^16, 42^ represents another key change allowing the full maturation of cells. However, this stage could not be included in our analyses since collection of neonatal tissue in human are extremely rare and ethically problematic. Nonetheless, our results established that the step-by-step differentiation of hepatocytes and cholangiocytes originates from a crosstalk among parenchymal and non-parenchymal cells. Of particular interest, stellate cells appear to have an underestimated importance in supporting hepatoblast self-renewal while building the hepatic niche. Such key developmental roles could be shared by many tissue specific fibroblasts involved in organ fibrosis^43, 44^ during chronic diseases. In addition, hepatoblasts/hepatocytes are likely to also influence the surrounding cells to establish their own niche and to direct their functional maturation through interactive feedback loops. Utilisation of this knowledge has enabled us to develop a culture system to grow hepatoblasts *in vitro* which provide a promising model system not only to study liver organogenesis, but also to produce cells for clinical applications. Nonetheless, it is important to note that further investigations are necessary to demonstrate the capacity of HBO to differentiate into fully functional hepatocytes and cholangiocytes after clonal isolation. Finally, our developmental map has revealed that differentiation of hPSCs diverge from a natural path of development at an early stage and then follow an *in vitro*-specific process. This divergence explains the fetal nature of cells generated by current protocols^45–50^ and suggests that improving the intermediate, specification steps of these protocols may be necessary to generate adult cells. Thus, understanding organ development remains the best approach for generating fully functional cell types *in vitro.* Our study illustrates how single-cell analyses can be combined with *in vitro* models to uncover the mechanisms driving the generation of functional hepatic cells *in vivo* and thus paves the way toward identifying factors for improving differentiation *in vitro*. This methodology and the resulting knowledge are likely to be transferable to other organs and will be useful for generating a diversity cell types for disease modelling and cell-based therapies.

## METHODS

### Adult human liver collection and dissociation

Liver samples were obtained under sterile conditions from deceased transplant organ donors as rapidly as possible after cessation of circulation. Tissue samples were transferred to the laboratory at 4°C in University of Wisconsin (UW) organ preservation solution. Biopsy tissue was taken from the patient, placed in room temperature HepatoZYME-SFM media and processed immediately. Protocols were developed from information in a previously published dissociation method^51^. The liver tissue was washed twice with warm DPBS with Ca^+2^Mg^+2^ + 0.5 mM EDTA to wash away blood and preservation medium, transferred into a petri dish and diced into small pieces (roughly 1 cm^2^ for resections and 0.25 cm^2^ for biopsies) using a scalpel. For large volumes of resected liver, the tissue was distributed evenly into four GentleMacs Tissue Dissociation C Tubes, whereas biopsy tissue was placed in a 1.5 ml Eppendorf tube. The tissue was washed twice with DPBS with Ca^+2^Mg^+2^, and then Liberase enzymatic digestion solution (0.2 Wünsch/ml) reconstituted in HepatoZYME media (without growth factors, which may influence the downstream scRNA-seq) containing DNAse I (2000 U/ml) was warmed to 37 °C for optimal enzymatic activity and added to each tube; 5 ml per tissue dissociator C tube and 1 ml per Eppendorf tube. The enzymatic digestion occurred in an incubating shaker at 37 °C and 200 RPM for 30 mins. The partially degraded extracellular tissue matrix was mechanically dissociated by running two “B” cycles using the “C tube” in the Miltenyi Biotec GentleMACS tissue dissociator. A 1:1 ratio of 20% FBS to 80% DPBS was added to neutralize the enzymatic reaction and the cell suspensions were filtered through 70 um filters to remove large pieces of debris, with large pieces gently mashed through the filter. Cells dissociated from donor resections were purified by centrifuging the cell suspensions at 50 x g, 4 °C for 5 mins to pellet the hepatocyte fraction, followed by centrifugation of the supernatant containing the non-parenchymal cell populations (NPCs) at 300 x g, 4 °C for 5 mins (1^st^ NPC fraction) and 650 x g, 4 °C for 7 mins (2^nd^ NPC fraction). Each hepatocyte pellet was resuspended in 5ml of DPBS, which was added to four separate cold 25% percoll solution. The density gradients layered by the cell suspension were centrifuged at 1250 x g, 4 °C for 20 mins without brake. Each purified hepatocyte pellet was resuspended in 5ml of red blood cell (RBC) lysis solution (Miltenyi Biotec 130-094-183) and incubated at room temperature for 10 mins. The cells were pelleted by centrifuging at 50 x g, 4 °C for 5 mins, after which the red supernatant containing the RBC lysate was aspirated. The hepatocytes were resuspended, pooled in 1ml of cold HepatoZYME-SFM, counted and stored on ice for downstream analyses. The two NPC fractions were pooled together in 3.1ml of DPBS, to which 900μl of debris removal solution is added. The Miltenyi Biotec Debris Removal (Miltenyi Biotec 130-109-398) protocol was followed according to manufacturer’s guidelines to yield a clean cell pellet. The cell pellets were then resuspended in 2 ml of RBC lysis buffer at room temperature and incubated at room temperature for 7-10 mins. The cells were pelleted by centrifuging at 650 x g, 4 °C for 5 mins. The red supernatant was aspirated and the NPC cell pellet was resuspended in 1ml of cold HepatoZYME media. The cells were counted, diluted using HepatoZYME media to use for single-cell transcriptomics, and stored on ice.

Non-parenchymal cell types were isolated from liver biopsies by first centrifuging the total cell suspension at 400 x g, 4 °C for 5 mins. The supernatant was aspirated and RBC lysis solution was added and incubated with the cells for 10 mins at room temperature. Subsequently, debris removal solution solution was used to clean remaining debris and dead cells according to manufacturer’s instructions. The clean pelleted cells were resuspended in HepatoZYME-SFM media at 4 °C, counted, and placed on ice.

### Immunohistochemistry of primary tissue sections

Primary human liver tissue was either snap-frozen in OCT, or fixed in 10% formalin and embedded in paraffin, sectioned onto slides, and stored for immunohistochemistry. Paraffin and OCT embedded samples were sectioned and placed onto slides for IHC by the tissue bank at Addenbrooke’s Hospital, Cambridge, UK. Paraffin embedded sections were deparaffinated and rehydrated by washing 3 times for 10 mins each with xylene followed by 3 washes for 5 mins each with ethanol. Frozen sections were brought to room temperature, followed by heat-induced antigen retrieval (HIER). All sections were washed for 5 mins in tap water, followed by 20 mins of HIER in 90 C citrate buffer with pH 6.0 and another 20 mins in the 90 C buffer at room temperature, allowing the buffer to cool ambiently. Sections were washed for 5 mins in tap water and 5 mins with DPBS + 0.1% TritonX-100, and then were blocked for 30 mins using DPBS + 10% donkey serum + 0.1% TritonX-100. Primary antibodies were diluted in DPBS + 1% DS and 0.1% Triton-X100 and incubated on the tissue sections overnight at 4 °C. The following day, the primary antibody was removed and the cells were washed 3 times for 5 mins each with DPBS + 0.1% TritonX-100. The secondary antibodies were diluted in 1:1,000 in DPBS + 1% DS and 0.1% Triton-X100 and incubated on the tissue in the dark at room temperature for 1 hr. The sections were washed 3 times for 5 mins each using DPBS, with the second wash containing 1:10,000 dilution of Hoechst counterstain. Fluorescence preservation mounting media and coverslips were placed on the tissue sections and stored at 4 °C prior to immunofluorescent imaging. Fluorescent IHC imaging was performed using the Zeiss LSM 700 confocal microscope and images were analysed using Imagej 1.48k software (Wayne Rasband, NIHR, USA, http://imagej.nih.gov/ij). All antibodies used are described in detail in Supplementary Table 22.

Hematoxylin and eosin (H&E), Masson’s trichrome, and other histological stains were performed by the histopathology and cytology department at Addenbrooke’s Hospital, Cambridge, UK. This department is jointly staffed by National Health Service (NHS) and University of Cambridge consultants with expertise in histopathology, and which regularly participates in internal and external quality control schemes. Staining was performed using Leica BOND-MAX automated IHC staining system. Histological imaging was performed using the Zeiss Axio Imager M2 and were processed using Zeiss Zen Blue software. All antibodies used are described in detail in Supplementary Table 22.

### Fetal human liver collection and dissociation

Primary human fetal tissue was obtained from patients undergoing elective terminations (REC- 96/085). The liver was dissected from the abdominal cavity and placed into a solution containing Hanks’ buffered saline solution (HBSS) supplemented with 1.07 Wünsch units/ml Liberase DH (Roche Applied Science) and 70 U/ml hyaluronidase (Sigma-Aldrich), and placed on a microplate shaker at 37 °C, 750 RPM, for 15 mins. The sample was subsequently washed three times in HBSS using centrifugation at 400 x g for 5 mins each. The single-cell suspension was then sorted for EPCAM/CD326 positive cells using CD326 microbeads (Miltenyi Biotec 130-061-101), according to the manufacturer’s guidelines.

### Establishment of hepatoblast organoids

The single cell suspension was resuspended in the hepatoblast organoid media (HBO-M); Advanced DMEM/F12 supplemented with HEPES, penicillin/streptomycin and glutamax, 2% B27, 20mM Nicotinamide, 2mM n-acetlycysteine, 50% WNT3A conditioned medium, 10% R-Spondin, 50ng/ml EGF, and 50uM A83-01. For long term culture (greater than 4 passages) 10uM Y27632 was required. To the resuspended cells was added a volume of Growth Factor Reduced Phenol Free Matrigel (Corning) to make the final solution up to 55% Matrigel by volume, and the mixture pipetted into 48 well plates (20uL per well). The plates were placed at 37°C for fifteen minutes to allow the mixture to set, and subsequently 200uL of fresh HBO- media applied to each well.

### Maintenance of hepatoblast organoid (HBO) cell lines

The culture medium (HBO-M, see above) was changed every 48-72 hours, and organoids mechanically passaged every 7-10 days. Organoids were passaged by scraping the gel away from the plate, pipetting the resulting solution into 1.5ml tubes, and pipetting the solution up and down to break individual organoids into pieces. If a precise cell number was required, the organoids can alternatively be processed to a single cell solution as described below, and then re-plated at the required dilution in the 55% Matrigel-medium solution.

### Dissociation of HBO to single cell solution

Organoids established in culture were dissociated to single cell suspensions for splitting/ single cell sequencing/ cell counting by first removing the media from each well and replacing with Cell Recovery Solution (Corning). The organoids in the Matrigel were then be scraped off the plate and placed on ice for 30 minutes to remove the Matrigel. The organoids were then washed with PBS and placed in TrypLE (ThermoFisher Scientific) for 15 mins at 37°C. Cells were finally washed three times in PBS or basal media.

### Establishment of biliary organoids (BO) and differentiated biliary organoids (DBO)

Biliary organoids were derived from samples taken from the livers of adult human deceased donors (National Research Ethics Committee East of England – Cambridge South 15/EE/0152). These were maintained and differentiated as per author’s guidelines^26, 52^.

### HBO differentiation to hepatocytes *in vitro*

Hepatocyte differentiation medium was made using complete HepatoZYME (Life Technologies 17705-021) (i.e. basal HepatoZYME supplemented with nonessential amino acids (ThermoFisher 11140050), chemically defined lipid concentrate (ThermoFisher 11905031), L-glutamine (2 mM), insulin (14 ng/ml), and transferrin (10 ng/ml). To this basal media was added either oncostatin-M (Sigma Aldrich) at a concentration of 20 ng/ml, EPO (R&D systems) at a concentration of 50 U/ml, or VEGFA (R&D systems) at a concentration of 50 ng/ml. This supplemented Hepatozyme was applied to HBO for seven days, applying fresh medium every 48 hours.

### HBO differentiation to cholangiocytes *in vitro*

HBO Maintenance medium was supplemented with TGF-β (2 ng/ml) and applied to established HBO lines for a total of seven days, applying fresh medium every 48 hours.

### Imaging of organoids

The medium was removed and replaced with 4% Paraformaldehyde solution for 20 minutes at room temperature to fix the organoids in situ, followed by three washes in PBS (each for five minutes). Fixed organoids could then be stored in PBS for later Immunohistochemistry. Immunostaining was performed by permeabilizing and blocking with 0.3% Triton X-100 (Sigma-Aldrich), and 10% donkey serum (Biorad) for three hours at room temperature, and then incubated overnight at 4 °C, with the primary antibody in 1% donkey serum. Samples were then washed with three one-hour PBS washes, followed by incubation with the fluorescent secondary antibody overnight at 4 °C. The samples were finally washed three times in PBS, with 1:10000 dilution Hoechst 33258 (ThermoFisher Scientific) added to the first wash.

### Generation of reporter lines

HBO were dissociated to a single cell suspension as described. Cells were transfected using a lentiviral vector containing tdTomato (rLV. EF1. tdTomato9, Clontech), according to the manufacturer’s guidelines. Following transfection HBO were grown in medium supplemented with puromycin (2 mg/ml) for ten days to select for stably transfected cells.

### Flow cytometry

Organoids were dissociated to single cell solution as described above. The single cells were then fixed and permeabilised in FixPerm (ThermoFisher Scientific) for 20 minutes, then washed three times in PBS before resuspension in PermWash (ThermoFisher Scientific). Blocking was performed using 10% donkey serum (Biorad) in PBS for 30 minutes. Primary antibodies were diluted 1:100 in 1% donkey serum in PBS and incubated with the cell suspension at room temperature for 1 hour. After three washes in PBS, cells were incubated with secondary antibodies diluted 1:1000 in 1% donkey serum in PBS. Data were collected using Cyan ADP flow cytometer, and analysed using FlowJo X (BD, FlowJo, LLC).

### Assessment of cell proliferation rates

Organoids were extracted and dissociated to single cells as described above. The cells were then counted using the countess II automated cell counter (Biorad), and the final number adjusted according to the number of wells used and the volume of resuspension.

### Cytochrome P450 enzyme activity

Cytochrome P450 enzymatic activity was assessed using Promega® P450-Glo^TM^ assay systems using Luciferin-IPA for and Luciferin-PFBE as surrogate markers for cytochrome P450 3A4 and cytochrome p450 3A5/7 respectively. Each substrate was diluted 1:1000 in freshly applied media as per manufacturer’s guidelines and placed on organoids in three-dimensional culture. After incubation at 37°C for four hours the media was collected and 50μl placed into a detection plate with 50μl detection reagent. The solution was then left for twenty minutes prior to being read via a luminometer. Readings were adjusted for the average of three control readings, with the control consisting of the medium and reagents that was kept at 37°C for four hours without contact with cells.

### Media protein analysis

Albumin, alpha-fetoprotein, apolipoprotein-B, and alpha-1-antitrypsin were detected in media by enzyme linked immunosorbent assay (performed by core biomedical assay laboratory, Cambridge University Hospitals). Concentrations were normalized to cell number.

### Murine blood protein analysis

Blood was drawn retro-orbitally for human albumin ELISA (Bethyl Laboratories) immediately prior to sacrifice at the termination of the experiment (27 days). Serum was separated by centrifugation and levels of human albumin were determined by an enzyme-linked immunosorbent assay (ELISA) using goat polyclonal capture and horseradish peroxidase– conjugated goat anti-human albumin detection antibodies (Bethyl Laboratories). Non-implanted FNRG mouse blood serum was included as a negative control.

### BODIPY assay

BODIPY 493/503 (ThermoFisher Scientific) was diluted 1:1000 in the organoid culture media and applied to organoids for 30 minutes. After this time the organoids were washed with fresh media and imaged in situ using a fluorescent microscope.

### Freezing/ thawing of HBO

Organoids were extracted from Matrigel using cell recovery solution as described above. The organoids were then washed in PBS and resuspended in Cell Culture Freezing Medium (ThermoFisher Scientific), placed in cryogenic vials (ThermoFisher Scientific), and cooled to -80°C using a “Mr. Frosty” Freezing Container (ThermoFisher Scientific). Organoids could then be kept long term in liquid nitrogen. To thaw, cryovials were kept at 37°C until the freezing medium had melted. Organoids were then washed three times in PBS to remove any residual freezing medium, and then re-plated as above.

### Culture of primary hepatocytes

Primary hepatocytes were purchased from Biopredic International (Rennes, France). Hepatocytes were isolated from human liver resects from three donors (two male and one female), and met the manufacturer’s quality control requirements in regards to viability, confluency and functionality, which was assessed by Phase I and II dependent enzymatic activity. Cells were purchased as monolayer cultures and maintained in William’s E (Gibco) supplemented with 1% Glutamine (Gibco), 1% Penicillin-streptomycin (Gibco), 700nM Insulin (Sigma) and 50µM Hydrocortisone (Sigma) for no longer than 48h upon receipt.

### Culture of HUVEC cell line for transplantation

Primary human umbilical vein endothelial cells (HUVECs, passage 2) were thawed and cultured in EGM-2 media (Lonza). Media was replaced every 48h. Cells were passaged once before implantation in mice.

### Encapsulation of HBO for implantation in mice

Hepatoblast organoids were isolated and then suspended in 55% matrigel (Corning) on ice. HUVECs were suspended in 10 mg/ml fibrin hydrogel (human thrombin, Sigma-Aldrich; bovine fibrinogen, Sigma-Aldrich) and mixed with organoids in un-polymerized matrigel at a 20:1 ratio. Cell/fibrin mix containing organoids and HUVECs was polymerized in disks from which 6 mm biopsies were punched to create “organoid tissues”. Each 6 mm implant contained approximately 3,500 organoids and 100,000 HUVECs.

### Implantation of HBO and induction of liver injury in mice

All surgical procedures were conducted according to protocols approved by the University of Washington Institutional Animal Care and Use Committees. 14 to 18-week-old female Fah−/− backcrossed to NOD, Rag2−/−, and Il2rg-null (FNRG) mice (Yecuris) were administered sustained release buprenorphrine and anesthetized with isofluorane. Three organoid tissues were sutured onto the inguinal fat pads of each mouse. Three mice received organoid tissues with Donor 4 hepatoblasts and 2 mice received organoid tissues with Donor 5 hepatoblasts. Incisions were closed aseptically. Nitisinone (NTBC) was withdrawn from animals’ drinking water immediately after implantation of organoid tissues and for 14 days after implantation to induce liver injury. NTBC was then reintroduced to the drinking water to allow for recovery, and then removed again after 4 days for the remainder of the experiment. Animals were sacrificed 27 days after implantation of organoid tissues.

### Immunostaining of harvested constructs from mice

Implants were harvested and fixed in 4% paraformaldehyde for 48 hours at 4°C. Excess fat was trimmed off of the implants, which were then dehydrated in graded ethanol (50-100%), embedded in paraffin, and sectioned using a microtome (6 mm). Some sections were histochemically stained with hematoxylin and eosin. For immunostaining, sections were blocked with normal donkey serum and incubated with primary antibodies against human (see supplementary table 4 for antibodies). To semi-quantify KRT19 distribution in nodules, graft nodules in which all cells were KRT18+/KRT19+ were tallied as “+”. Nodules with both KRT18+/KRT19+ and KRT18+/KRT19- cells were tallied as “+/-“. Nodules with only KRT18+/KRT19- cells were tallied as “-“. Nodules in each category were summed across each tissue section and divided by total KRT18+ grafts in the section to acquire percentages in each animal, with each data point representing one animal.

### Fluorescent *in situ* mRNA hybridization of primary fetal tissue

Primary fetal human liver tissue was fixed in 10% formalin and embedded in paraffin (FFPE), sectioned on to slides and stored for smFISH. FFPE slides were baked at 65°c for 1 hour. Slides were deparaffinized with two washes for 10mins each in xylene solution (Bond™ Dewax Solution, Leica AR9222) and two washes for 5 mins each in PBS(1x). Slides were air dried before automated smFISH protocol.

Multiplex smFISH was performed on a Leica BondRX fully automated stainer, using RNAScope© Multiplex Fluorescent V2 technology (Advanced Cell Diagnostics 322000). Slides underwent heat-induced epitope retrieval with Epitope Retrieval Solution 2 (pH 9.0, Leica AR9640) at 95°C for 5 mins. Slides were then incubated in RNAScope© Protease III reagent (ACD 322340) at 42°C for 15 mins, before being treated with RNAScope© Hydrogen Peroxide (ACD 322330) for 10 mins at RT to inactivate endogenous peroxidases.

All double-Z mRNA probes were designed by ACD for RNAScope on Leica Automated Systems. Slides were incubated in RNAScope 2.5 LS probes (designed against human genes *Rspo3, Lgr5, Alb, Pdgfrβ, Lrp5, Des, Dkk1, Epcam, Notch2, Kdr, Dll4, Wnt2b* and *Wnt4)* for 2 hours at RT. DNA amplification trees were built through consecutive incubations in AMP1 (preamplifier, ACD 323101), AMP2 (background reduction, ACD 323102) and AMP3 (amplifier, ACD 323103) reagents for 15 to 30 mins each at 42°C. Slides were washed in LS Rinse buffer (ACD 320058) between incubations.

After amplification, probe channels were detected sequentially via HRP–TSA labelling. To develop the C1–C3 probe signals, samples were incubated in channel-specific horseradish peroxidase (HRP) reagents for 30 mins, TSA fluorophores for 30 min and HRP-blocking reagent for 15 min at 42 °C. The probes in C1, C2 and C3 channels were labelled using Opal 520 (Akoya FP1487001KT), Opal 570 (Akoya FP1488001KT), and Opal 650 (Akoya FP1496001KT) fluorophores (diluted 1:500) respectively. The C4 probe complexes were first incubated with TSA–Biotin (Akoya NEL700A001KT, 1:250) for 30 min at RT, followed by streptavidin-conjugated Atto425 (Sigma 56759, 1:400) for 30 min at RT. Samples were then incubated in DAPI (Sigma, 0.25µg /ml) for 20 mins at RT, to mark cell nuclei. Slides were briefly air-dried and manually mounted using ∼90 µl of Prolong Diamond Antifade (Fisher Scientific) and standard coverslips (24 × 50 mm^2^; Fisher Scientific). Slides were dried at RT for 24 hrs before storage at 4°C for >24 hrs before imaging.

SmFISH stained fetal liver slides were imaged on an Operetta CLS high-content screening microscope (Perkin Elmer). Image acquisition was controlled using Perkin Elmer’s Harmony software. High resolution smFISH images were acquired in confocal mode using an sCMOS camera and x40 NA 1.1 automated water-dispensing objective. Each field and channel were imaged with a z-stack of 20 planes with a 1µm step size between planes. All appropriate fields of the tissue section were manually selected and imaged with an 8% overlap.

### HBO/stellate cells co-cultures

To model hepatic stellate cells *in vitro* we used the LX2 cell line, which was modified to obtain a CFP reporter cell line (rLV.EF1.AmCyan1-9 [0039VCT] Clontech, Takara Bio Europe). The LX2 cell line was maintained in DMEM, high glucose, GlutaMAX™ Supplement (Thermo Fisher Scientific, 10566016) in presence of 5% FBS (SLS, SH30070.03) and 1% Pen/Strep. When both HBO and LX2 cultures reached confluence for co-cultures, HBO were manually scraped and cells incubated on ice for 5minutes/well (for a total of 20minutes), while LX2 were treated with Trypsin-EDTA (0.05%) (Thermo Fisher Scientific, 25300054) for 5minutes at 37C. Cells were spun down at 600g and re-suspended in 1ml of medium and counted. Cells were then pooled together at the following ratio: 70%HBO / 30%LX2, for a maximum of 20’000 cells/well. Matrigel was added to the cell suspension, and cells were seeded as described for HBO cultures, in complete HBO medium. Treatments of WNT or RSPO withdrawal started 2 days after seeding, and cells were cultured for 7 days.

### Generation and culture of hPSC lines

The human pluripotent stem cell (hPSC) lines used in this study were generated previously in our lab and collaborators’ labs. Cells were maintained in defined culture described previously^32, 36, 45^, using Essential 8 media with the provided nutrient supplement (ThermoFisher A1517001) or DMEM/F-12 basal media (ThermoFisher 11320033) supplemented with insulin-transferring-selenium (2x) (ThermoFisher 41400045), sodium bicarbonate (1.74 g/L) (ThermoFisher 25080094), L-ascorbic acid 2-phosphate, hFGF2 (25 ng/ml) and TGFβ (2 ng/ml) (see Supplementary Table 19 for all media compositions). The cells were incubated at 37 C and 5% CO2 with media changed daily. Typically, cells were maintained for up to one week before passaging. To passage the cells, DPBS was used to wash each well being passaged, followed by incubation at 37 °C with 0.5 mM EDTA in DPBS for 4 mins. The 0.5mM EDTA in DPBS was aspirated, fresh E8 media was added, and a pipette was used to flush the wells to detach the cells and break them into 5-10 cell clumps. The cells were transferred into a tube and allowed to settle, thus eliminating the small clumps and single cells.

The supernatant was aspirated, the pellet was resuspended in complete E8 media and plated, typically in a 1:12 ratio split.

### Differentiation of hPSCs into hepatocyte-like cells (HLC)

The differentiation of hiPSCs to HLCs followed the protocol previously published by the Vallier lab^45, 46^. Prior to differentiation, 12-well plates were coated with 0.1% gelatin for an hour at 37 °C followed by MEF medium overnight at 37 °C and washed with DPBS before use. The hiPSC were passaged at 70-90% confluency by washing with DPBS and then adding Accutase cell dissociation reagent (ThermoFisher A1110501) and incubating at 37 °C for 4 mins. After incubation, accutase was diluted at a 1:1 ratio with complete E8 media (Gibco A1517001) and the supernatant was pipetted gently to mechanically break any clumps into single cells. The resulting suspension was centrifuged at ∼350 x g for 3 mins to pellet, resuspended in E8 media and counted. The hiPSCs were plated at a density of 50,000-60,000 cells/cm^2^ in 12-well plates in complete E8 media supplemented with Y27632 ROCK inhibitor (10 uM) on 0.1% gelatin/MEF-coated plates. The medium was changed the following day to complete E8 without ROCK inhibitor. On day 1 of differentiation (2 days after plating), CDM-PVA media supplemented with activin (100 ng/ml), FGF2 (80 ng/ml), BMP4 (10 ng/ml), LY294002 (10 uM), and CHIR99021 (3 uM) was added to the cells to induce endoderm formation (see Supplementary Table 24 for all complete media compositions). CHIR99021 was removed from this medium on day 2 of differentiation. On day 3, RPMI-1640 with nonessential amino acids (ThermoFisher 11140050) and B27 supplement (ThermoFisher 17504044) was supplemented with activin (100 ng/ml) and FGF2 (80 ng/ml). Foregut differentiation was initiated on day 4 and carried out until day 8 by changing media to complete RPMI media supplemented with activin (50 ng/ml). The hepatoblast and subsequent hepatocyte phenotype was induced by changing medium to HepatoZYME-SFM (Life Technologies 17705-021) supplemented with nonessential amino acids (ThermoFisher 11140050), chemically defined lipid concentrate (ThermoFisher 11905031), L-glutamine (2 mM), insulin (14 ng/ml), transferrin (10 ng/ml), oncostatin M (OSM) (20 ng/ml) and hepatocyte growth factor (HGF) (50 ng/ml) from day 9 to day 33.

### Differentiation of hPSCs into cholangiocyte-like cells (CLC)

Differentiation of hPSCs to CLCs followed the protocol previously published in our lab^36^. Prior to differentiation, 12-well plates were coated with 0.1% gelatin for an hour at 37 °C followed by MEF medium overnight at 37 °C and washed with DPBS before use. The hiPSC were passaged at 70-90% confluency by washing with DPBS and then adding Accutase cell dissociation reagent (ThermoFisher A1110501) and incubating at 37 °C for 4 mins. After incubation, accutase was diluted at a 1:1 ratio with complete E8 media (Gibco A1517001) and the supernatant was pipetted gently to mechanically break any clumps into single cells. The resulting suspension was centrifuged at ∼350 x g for 3 mins to pellet, resuspended in E8 medium and counted. The hPSC were plated in complete E8 medium supplemented with ROCK inhibitor Y27632 (10 uM) at a density of 50,000-60,000 cells/cm^2^ on 0.1% gelatin/MEF-coated plates. The media was changed the following day to E8 without ROCK inhibitor, and two days after plating, the differentiation was started by changing the media to CDM-PVA supplemented with activin (100 ng/ml), FGF2 (80 ng/ml), BMP4 (10 ng/ml), LY294002 (10 uM), and CHIR99021 (3 uM) to induce endoderm formation. The same medium was used the following day (day 2) without CHIR99021. On day 3, RPMI-1640 with nonessential amino acids and B27 supplement (ThermoFisher 17504044) (RPMI+ media) was supplemented with activin (100 ng/ml) and FGF2 (80 ng/ml) only. Hepatoblasts were induced from day 9 to day 12 using RPMI+ medium supplemented with SB (10 uM) and BMP4 (50 ng/ml). This bipotent progenitor was directed toward the cholangiocyte lineage from day 13 to day 16 by feeding with RPMI+ media supplemented with retinoic acid (3 uM), FGF10 (50 ng/ml), and activin (50 ng/ml). The mature cholangiocyte phenotype was induced by re-plating the cells in 3D culture using a 1:2 ratio of cell suspension to Matrigel. The cells matured from day 17 to day 26 in 3D culture with common bile duct media (CBD media) supplemented with EGF (50 ng/ml) and forskolin (10 uM), yielding CLCs at day 26 of differentiation.

### Differentiation of hPSC into hepatic stellate-like cells (HSLC)

This differentiation protocol was gathered from a previously published paper^22^. Prior to differentiation, 12-well plates coated with a 1:50 ratio of reduced growth factor Matrigel and low glucose DMEM medium overnight at 37 °C and washed with DPBS before use. The hiPSC were passaged at 70-90% confluency by washing with DPBS and then adding Accutase cell dissociation reagent (ThermoFisher A1110501) and incubating at 37 °C for 4 mins. After incubation, accutase was diluted at a 1:1 ratio with complete E8 media (Gibco A1517001) and the supernatant was pipetted gently to mechanically break any clumps into single cells. The resulting suspension was centrifuged at ∼350 x g for 3 mins to pellet the cells, which were then resuspended in E8 medium and counted. The hiPSC were plated in complete E8 media supplemented with Y27632 (10 uM) on 12-well plates coated with a 1:50 ratio of reduced growth factor Matrigel and low glucose DMEM overnight at a concentration of 90,000 cells/cm^2^. The media was changed to E8 without ROCK inhibitor the following day. The cells were differentiated to mesoderm by adding DMEM-MCDB 201 media supplemented with BMP4 (20 ng/ml) on day 1 and day 3 of differentiation. On day 5, a mesenchymal phenotype was induced by adding DMEM-MCDB 201 media with BMP4 (20 ng/ml), FGF1 (20 ng/ml), and FGF3 (20 ng/ml). These cells transitioned to liver mesothelium by adding DMEM-MCDB 201 supplemented with retinoic acid (RA) (5 uM), palmitic acid (PA) (100 uM), FGF1 (20 ng/ml), and FGF3 (20 ng/ml) on day 7. From days 9 to 13, the cells were fed every 2 days with RA (5 uM) and PA (100 uM) to attain a fetal hepatic stellate-like cell phenotype (HSLC).

### Differentiation of hPSC into endothelial-like cells (ELC)

This protocol was adapted from previously published papers^21, 37^. Prior to differentiation, 12- well plates were coated with 0.1% gelatin for an hour at 37 °C followed by MEF media overnight at 37 °C and washed with DPBS before use. The hiPSC were passaged at 70-90% confluency by washing with DPBS and then adding Accutase cell dissociation reagent (ThermoFisher A1110501) and incubating at 37 °C for 4 mins. After incubation, accutase was diluted at a 1:1 ratio with complete E8 media (Gibco A1517001) and the supernatant was pipetted gently to mechanically break any clumps into single cells. The resulting suspension was centrifuged at ∼350 x g for 3 mins to pellet, resuspended in E8 media and counted. The hiPSC were plated in E8 media supplemented with Y27632 (10 uM) at a density of 45,000 cells/cm^2^ on 0.1% gelatin/MEF-coated plates. Differentiation was begun the follow day (day 1) by inducing mesoderm using CDM-PVA media supplemented with FGF2 (20 ng/ml), BMP4 (10 ng/ml), and LY294002 (10 uM). On day 2.5, the cells were fed with Stempro-34 media with VEGFA (200 ng/ml), forskolin (2 uM), and L-ascorbic acid (1 mM). The media was changed every day until day 5.5, after which the fetal endothelial-like cells (ELCs) were used for downstream analyses or re-plating.

### Differentiation of hiPSC into macrophage-like cells (MLC)

This protocol was adapted from a previously published protocol^38^. The hiPSC were passaged at 70-90% confluency and plated onto an ultra-low adherence 96-well plate with embryoid body (EB) media consisting of E8 supplemented with Y27632 (10 uM), BMP4 (50 ng/ml), SCF (20 ng/ml), and VEGF (50 ng/ml). The cells were incubated for 4 days with half of the media in each well being replaced with fresh media after 2 days. On day 4, the EBs were transferred to a 6-well plate coated with 0.1% gelatin in DPBS, and X-VIVO 15 media supplemented with Glutamax (2 mM), 2-mercaptoethanol (55 uM), M-CSF (100 ng/ml), and IL-3 (25 ng/ml) was added to the wells. Every 5 days, for roughly 10 days, 2/3 of the media in each well was changed. On day 14, the EBs began production of macrophage progenitors. These floating progenitors were collected with the supernatant and plated on uncoated dishes in RPMI + 10% FBS supplemented with M-CSF (100 ng/ml) On day 7 after macrophage progenitor plating, the differentiated macrophages were collected for downstream analyses.

### Differentiation of hiPSC into stellate-endothelial progenitor cells (SEpro)

Cells were plated and differentiated according to the endothelial-like cell (ELC) differentiation protocol described previously. At day 3.5 of this protocol, the SEpro cells were present in culture and collected to analyses. To test the bipotentiality of these cells, they were dissociated from the palte into a single-cell suspensions by incubating with TryplE for 20 min at 37 °C, resuspending in culture media and replating at a density of 100,000 cells/cm^2^. These cells were subjected to the hepatic stellate-like cell (HSLC) differentiation conditions beginning at day 7 of this protocol and continuing for at least 4 days to produce HSLCs.

### Lentivirus production

HEK 293T cells were expanded and subsequently passaged by washing with DPBS and incubating with 0.5% EDTA in DPBS. The cells were pelleted, resuspended in MEF media, counted, and plated in T150 flasks at a density of 1.8*107 cells per flask. The cells were allowed to attach for 24 hr at 37 °C before transfection with the three lentiviral (LV) plasmids. In 15ml Falcon tubes (one per virus being produced), 25 ug of pWPT-transgene vector cloned from the pWPT-GFP vector (addgene #12255), 14 ug of psPAX2 LV helper plasmid (addgene #12260), and 6 ug of pMD2.G LV packaging plasmid (addgene #12259) were added to room temperature basal RPMI media. The solution was vortexed, 68 ul of TurboFect transfection reagent (ThermoFisher R0532) was added, and complexes were formed for 30 min at room temperature. The complexes were gently pipetted into the T150 flasks containing the HEK cells, one flask per LV transfection, and incubated for 18-24 hr at 37 °C. Following the 18-24 hr transfection period, the cells were washed with DPBS and 15 ml of MEF media was added. After 24-36 hr, the supernatant was harvested and filtered through 0.45 μm filters. DNaseI (5 U/ml) and MgCl2 (1 mM) was added (optional due to the presence of Mg already in the supernatant media), and incubated at room temperature for 20 min. Clontech Lenti-X Concentration (Takara 631232) solution was used to concentration the LV batches by adding 5.0 ml of Lenti-X Concentrator to 15.0 ml of supernatant (1:3 ratio), and incubating at 4 °C for 30 min to 12 hr. The solution was centrifuged at 1500 x g, 4 °C for 45 min, after which the supernatant was aspirated, and the pellet was resuspended in 250 ul (1:100 of the pre concentrated LV containing supernatant volume). The LV batches were frozen at -80 °C in single-use aliquots for experiments to prevent drastic decreases in efficacy caused by free-thaw cycles.

### Lentiviral transduction of iPSC

Before transduction, fresh hepatocyte-like cell differentiation media, HepatoZYME-SFM (see Supplementary Table 24 for media compositions), was added to cells in a 12-well plate format. Lentiviral aliquots were thawed on ice and added to the desired treatment wells under sterile conditions. Polybrene (PB) was added to a final concentration of 10 ug/ml to each of the wells, and the plates were rocked gently to ensure even distribution of lentivirus on the adherent cells. The cells were incubated at 37 °C for 24 hrs to allow the lentivirus to infect the cells and stably integrate the transgene into the host genome. After 24 hrs, the cells were washed with DPBS and fresh HepatoZYME-SFM with OSM (20 ng/ml) and HGF (50 ng/ml) was added to the cells. This washing procedure was repeated again at 48 hrs and 72 hrs post-transduction to ensure the removal of any remaining lentivirus. In these experiments, hiPSC-derived hepatocyte-like cells were transduced on day 15 of differentiation and assayed on day 23/25, or transduced on day 25 of differentiation and assayed on day 33.

### RNA extraction, cDNA reverse transcription, and quantitative real-time PCR

Cellular RNA was extracted using the Sigma Aldrich GenElute Total RNA Purification Kit (RNB100-100RXN) following the manufacturer’s suggested protocol, including the optional DNAse incubation step to increase RNA purity. The cDNA was produced using previously established reverse transcription methods. RNA quantity and purity was analysed, and 500 ng of RNA was added to each 11.8 ul reaction along with 0.5 ul of random primer (Promega C1181) and 1 ul of dNTP (Promega U1511). The remaining volume was brought up to 11.8 ul using nuclease-free water (ThermoFisher AM9937). The reactions were incubated at 65 °C for 5 mins to denature the RNA and random primers, then snap cooled on ice and placed at room temperature. A master mix of 4 ul of 5x 1^st^ strand buffer (ThermoFisher 18064-071), 2 ul of 0.1M DTT, 0.5 ul RNase OUT (ThermoFisher 10777-019), and 0.2 ul of Superscript II (ThermoFisher 18064-071) was prepared per reaction, and scaled based on the number of reactions in the experiment. The master mix was vortexed and 6.7 ul of master mix was added to each reaction. The reactions were then incubated on a heat block at 25 °C for 10 mins, 42 °C for 50 mins, and lastly 70 °C for 15 mins. The final cDNA was diluted to 600 ul using nuclease-free water.

Reverse transcription was followed by qPCR in a 384-well format using biological triplicates and technical duplicates in each experiment. All primer pairs are diluted to 100 uM stocks and subsequently diluted to 5 uM for use in qPCR. The reaction volumes in a 384-well non-skirted plate (ThermoFisher AB-0600) was 10 ul consisting of 0.4 of ul forward primer (Sigma Aldrich), 0.4 ul of reverse primer (Sigma Aldrich), 5.0 ul of Kapa Syber Fast Low Rox (Sigma Aldrich KK4622), 1.2 ul of nuclease-free water, and 3 ul of cDNA. The plates were sealed with optically clear adhesion covers (ThermoFisher 4360954) and run using the following cycling parameters: 95 °C for 3 mins, 95 °C for 5 secs, 60 °C for 30 secs, cycle to step 2 another 39 times, and lastly melt curve from 65 °C to 95 °C. All expression values (2^- ΔCt^) were normalized to housekeeping genes PBGD and RPLP0. All primers used are described in detail in Supplementary Table 23.

### Single-cell encapsulation, library preparation, sequencing, and data processing

Single cell-suspensions from primary tissue and *in vitro* culture were loaded onto the Chromium controller by 10X Genomics, which is a droplet-based single-cell capture platform. The individual cells flowed through the microfluidic chip, were lysed and tagged by a bead containing unique molecular identifiers (UMIs) and were encapsulated in an oil droplet. This resulting emulsion was amplified through reverse-transcription, and follows library preparation as dictated by the 10X Genomics manual. The resulting libraries were sequenced on the Illumina HiSeq 4000 platform. These files were aligned to the GRCh38 human genome and pre-processed using the CellRanger 10X Genomics software for downstream analyses.

### Reads mapping

The 10X single cell sequencing data were mapped with the Cellranger program (version 2.0.2)^53^ to the reference ‘refdata-cellranger-GRCh38-1.2.0’, which is the human genome ‘GRCh38’ downloaded from (https://support.10xgenomics.com/single-cell-gene-expression/software/downloads/latest). Reads for each sample were first imported into an ‘AnnDatà object (https://anndata.readthedocs.io/en/latest/ index.html) in the SCANPY^54^ framework, count matrices were then concatenated into a total matrix of 273,033 cells and 33,694 genes.

### Quality control

Cells expressing fewer than 1000 counts, fewer than 500 genes or more than 40% mitochondrial content were excluded. Application of such filter selected a total number of 237978 cells, which is 87% of the raw number of cells. Genes expressed in fewer than 3 cells were filtered out, leaving 29907 genes (89% of the total number of genes). Doublets were identified by applying two doublet prediction methods: Doubletdetection^55^ and Scrublet^56^.

### Normalisation and regression

Read counts were log-normalised in the SCANPY framework using total count normalisation (scaling factor 10,000). Highly variable genes were selected for downstream analysis based on their normalised dispersion, obtained by scaling with the mean and standard deviation of the dispersions for genes falling into a given bin for mean expression of genes (‘highly_variable_genes’ routine in SCANPY). Technical variation was regressed out using a linear regression model (‘regress_out’ function from the python package of ‘NaiveDÈ^57^), which allows the user to specify wanted variance when cell types or stages are known. Alternatively, the ‘COMBAT’ function from the svaseq^58^ package could have been equivalently used. Regressed values were scaled by maintaining the default maximum value of 10 for each gene expression in each cell (‘scalè routine in SCANPY).

### Clustering, annotation and dimensionality reduction

Clusters were detected by applying the graph-based, community detection method ‘Louvain’. Annotation of clusters was based on the expression of known markers. Since Louvain clustering does not necessarily correlate with biologically meaningful clusters and can over-cluster cells associated with the same cell type, clusters with the same annotation were merged. Principal Component Analysis (PCA), t-Distributed Stochastic Neighbor Embedding (TSNE)^59^ and Uniform Manifold Approximation and Projection (UMAP)^60^ plots were calculated with the SCANPY routines.

### Graph-abstraction

Transcriptional similarity among stages was quantified and visualised using the method: Partition-Based Graph Abstraction (PAGA)^61^. PAGA generates a graph-like map of cells based on estimated connectivities which preserves both the continuous and disconnected structure of the data at multiple resolutions. In a PAGA graph, nodes correspond to cell groups and edge weights quantify the connectivity between groups. Such weights represent the confidence in the presence of an actual connection by allowing discarding spurious, noise-related connections. The highest the connectivity values, the highest the confidence of a connection. In our PAGA graphs, we used connectivity estimates to analyse the relative similarity among stages.

### Pseudotemporal ordering and alignment

Time-related genes were selected as markers in collection time (Wilcoxon-Rank-Sum test, z-score>10). Diffusion pseudotime of time-related genes was derived using the DPT^54^ routine implemented in SCANPY. Comparison of pseudotemporal trajectories was performed within the cellAlign^62^ framework. cellAlign applies dynamic time warping to compare the dynamics of two single-cell trajectories using a common gene set and to identify local areas of highly conserved expression. The algorithm calculates pairwise distances between ordered points along the two trajectories in gene expression space. cellAlign then finds an optimal path through the matrix of pairwise distances which preserves pseudotemporal ordering and minimises the overall distance between the matched cells. We applied cellAlign onto all corresponding pairs of hiPSCs and primary cell types by selecting genes used for calculating diffusion pseudotime both in hiPSCs and primary cell types. Cells whose distances were lower than a 0.25 quantile threshold were annotated as aligned in Figure 5a.

### CellPhoneDB analysis

The significance of cellular interactions between cell types was calculated with a publicly available repository of curated receptors, ligands and their interactions (CellPhoneDB^63^, v2.0). Normalised data for primary tissues at each collection time point were used as input. Significance of interactions was calculated based on random permutations of cluster labels to generate a null distribution. Interactions were considered significant based on the default p-value threshold (p-value<0.05).

### Analysis of published RNA-sequencing data

Bulk RNA-Seq counts for differentiated biliary organoids (DBO), fetal hepatocyte organoids (FHO), primary adult hepatocytes (PAH) and primary fetal hepatocytes (PFH) were downloaded from GEO (series GSE111301, samples GSM3490461, GSM3490463, GSM3490466, GSM3490467, GSM3490468, GSM3490470). Similarly to *Hu et. al. 2018*, counts were normalised using the DESeq2 package in the R environment by applying the variance stabilising transformation. Cell Ranger feature-barcode matrices derived from hepatoblast organoids (HBO) were filtered, based on number of genes detected and percentage of reads that map to the mitochondrial genome (<0.1), log-normalised and averaged using the Seurat package. PCA analysis of the merged bulk and averaged single cell samples was performed using the prcomp package in R.

### Statistics and reproducibility

Primary cell-specific stage-identification was determined based on the standard Louvain clustering routine of scRNA-seq data in SCANPY. Calculation of differentially expressed genes was calculated using a p-value threshold (p<0.01) and absolute log-fold change threshold (|log-fold change| > 1.0). Calculation of time-related genes were selected based on z-score (z-score>10) thresholds for statistical significance using the Wilcoxon-Rank-Sum test, as stated above. Diffusion pseudotime was calculated using the SCANPY computational routine. Significant values for CellPhoneDB ligand-receptor interactions were selected based on a p-value threshold for significance (p-value<0.05). Traditional ORA pathway enrichment of the interactions revealed by CellPhoneDB were selected for statistical significance based on a p-value threshold (p<1e-16). Transcriptomic shifts between control hPSC-derived cells and control, untreated cells were determined based on Louvain clustering with standard parameters in SCANPY, as well as through differential gene expression based on a p-value threshold (p<0.01) and absolute log-fold change threshold (|log-fold change| > 1.0).

Statistical tests comparing groups in qPCR analyses were calculated using GraphPad Prism. Unpaired samples were compared for each condition using unpaired, two-tailed t-tests, as annotated in the corresponding figure legend (*p<0.05, **p<0.01, ***p<0.001, and **** p<0.0001, error bars denoting SEM). All immunofluorescent and histology stains are representative and correlate to sequencing results. All biologically-independent replicates are stated explicitly in their respective figure legends.

### Ethical approval

Primary human adult liver samples were obtained from deceased transplant organ donors (National Research Ethics Committee East of England – Cambridge South 15/EE/0152). Primary human adult liver biopsies were collected from living patients under ethical consent to be used in research (North West - Preston Research Ethics Committee 14/NW/1146). Primary human fetal liver tissue was obtained from patients undergoing elective terminations (REC-96/085). All human tissue was used after obtaining informed consent for use in research.

## Data availability

All raw single-cell RNA sequencing data has been deposited to ArrayExpress under accession E-MTAB-8210. Fetal liver sequencing data has been deposited to ArrayExpress under accession E-MTAB-7189. Fetal liver sequencing data from Popescu et al. has been deposited to ArrayExpress under accession E-MTAB-7407. Adult liver sequencing data from MacParland et al. and Ramachandran et al. have been deposited to NCBI GEO under accession GSE115469 and GSE136103, respectively. All data sources are described in Supplementary Table 1. Any additional data is available upon reasonable request.

## Code availability

All Python and R scripts supporting the findings of this paper are available upon reasonable requests. CellPhoneDB is available at www.CellPhoneDB.org.

## Acknowledgements

With thanks to Komal Nayak (University Department of Paediatrics, Cambridge) for help with maintenance of cell lines and technical support. The Cellular Genetics department at Cambridge University Hospital (Dr Ingrid Simonic) for performing comparative genomic hybridisation, the Core Biochemical Assay Laboratory at Cambridge University Hospitals (Keith Burling) for ELISA analysis of culture media, Roger Barker and Xiaoling He (John Van Geest Centre for Brain Repair, University of Cambridge) for their help accessing tissue, Fredrik Johansson (University of Washington) for surgical and animal support, and the Tietze Foundation for funding support. We acknowledge the Cambridge Biorepository for Translational Medicine for the provision of human tissue used in the study. We acknowledge the NHS Addenbrooke’s Hospital Tissue Bank for sectioning samples for histology and the Histopathology and Cytology service for immunohistochemistry staining of primary liver tissue. We thank the Cambridge Stem Cell Institute and the Imaging facility. B.W. was supported by the Gates Cambridge funding program. A.R. was supported by Wellcome Translational Medicine and Therapeutics Clinician PhD programme. The L.V. lab and C.M.M iare funded by the Chan Zuckerberg Initiative project grant: “a reference cell atlas of human liver diversity over a lifespan”, the Cambridge University Hospitals National Institute for Health Biomedical Research Centre and the core support grant from the Wellcome Trust and Medical Research Council of the Wellcome–Medical Research Council Cambridge Stem Cell Institute. C.A. and D.G received funding from the Open Targets consortium (OTAR026 project) and the Wellcome Sanger core funding (WT206194) A.W.J., F.S., L.V., A.E.M. and K.S.-P. gratefully acknowledge support from the Rosetrees Trust (REAG/240 and NMZG/233).

## Author contributions

B.W. performed experimental design, data generation of primary human liver and hPSC differentiation single-cell RNA sequencing, tissue processing and analyses, data analysis and biological interpretation, and manuscript preparation; A.R. performed experimental design, generation of organoid systems and their differentiation, data generation, tissue dissection, processing, and analyses, data analysis and biological interpretation, and manuscript preparation; D.M. performed computational analyses and interpretation; S.S. and K.S. conceived of and performed the *in vivo* experiments, contributed to manuscript preparation, and generated figures; J.K. performed experiments and provided analytical support and supervision. Z.M. performed computational analyses and interpretation; R.A.T. performed experimental design, tissue processing, critically appraised the manuscript, and provided intellectual contributions; C.M.M. performed tissue dissociation protocol development, immunofluorescence analysis and provided intellectual contributions; K.M. provided primary adult human liver tissue directly from deceased donors in clinic; J.G.B. provided technical support while experiments were planned and performed; R.B. provided published data to include in the study and critically appraised the manuscript with assistance from E.S. and D.M.P.; C.A. provided data and intellectual contributions with advice from D.G.; S.M. provided data and intellectual contributions with G.B.; D.O. provided intellectual contributions, flow cytometry analysis, and critically appraised the manuscript; E.D.Z. contributed to in vitro experiments; F.S. performed tissue processing, provided data, provided intellectual contributions, and critically appraised the manuscript; O.C.T. and K.S.P performed direct hepatic and renal mouse injections, K.S.P. additionally provided primary human liver tissue and intellectual contributions; M.H. provided published datasets, intellectual contributions, and critically appraised the manuscript; M.Z. supervised organoid generation and provided intellectual contributions, S.T. performed computational analyses planning, data interpretation, provided intellectual contributions, and critically appraised the manuscript; L.V. conceived the study, performed experimental design, interpreted data and analyses, and prepared the manuscript.

---

**Extended Data Fig. 1.**
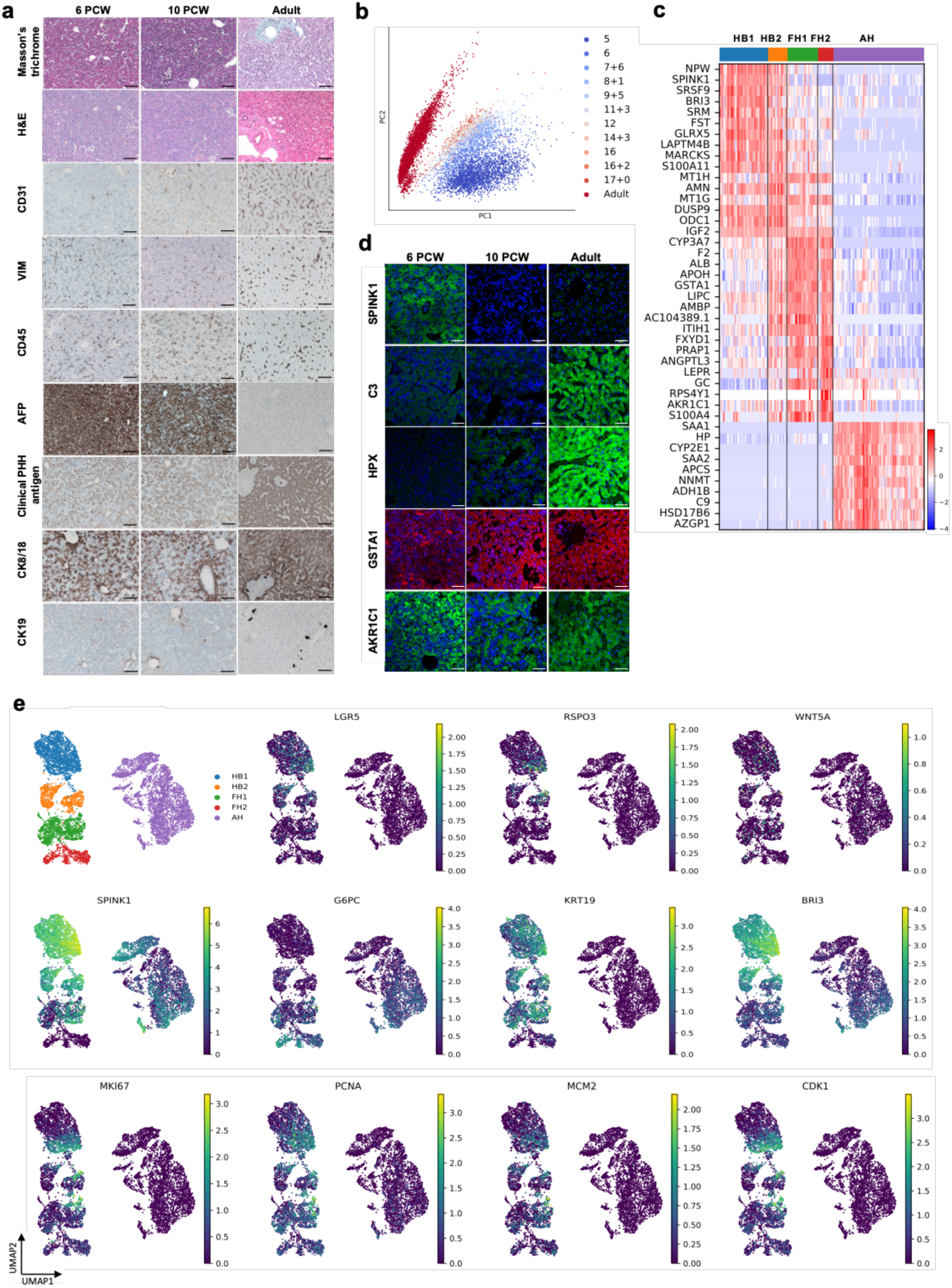
Characterisation of the proliferation and differentiation of cells within the developing human liver. **a,** Histology sections of key transcriptional and morphological events of primary fetal and adult liver. H&E and histology sections of primary fetal and adult liver spanning the key developmental ages within this study. From early fetal (6 PCW ± 4-5 days), intermediate fetal (10 PCW) and adult developmental stages. These immunostainings show transcriptional and morphological events including loss of AFP and increase of hepatocyte-specific antigen during development. Induction of CK19 as cholangiocyte are specified, with noticeable increased expression around duct formation in the 10 PCW sample. Note the increase of CK8/18 expressing cells as the liver develops, and the necessary increase in vasculature shown by CD31 expression. Importantly, VIM and CD45, staining for mesenchymal (stellate) cells and immune (Kupffer) cells respectively, are present throughout all stages of liver development. Scale bars = 100 um. **b,** PCA of fetal hepatoblasts/hepatocytes from 5-17 PCW and adult hepatocytes. **c,** Heatmap of the top 10 time-related differentially expressed genes (DEG) at each primary human hepatocyte developmental stage (Wilcoxon-Rank-Sum test, z-score>10). HB1, hepatoblast stage 1; HB2, hepatoblast stage 2; FH1, fetal hepatocyte stage 1; FH2, fetal hepatocyte stage 2; AH, adult hepatocyte. **d,** IHC analyses showing the expression of stage-specific markers on liver tissue sections. Scale bars = 50 um. HB, sample dated between 5-7 PCW; FH, sample dated at 11 PCW; AH, adult liver. **e,** UMAP representation of hepatoblast/fetal hepatocyte markers (SPINK1, G6PC, and BRI3) and WNT pathway markers (WNT5A, LGR5 and RSPO3). UMAP visualization also shows cell proliferation markers and cell cycle regulators across the hepatocyte developmental stages, showing progressive loss of proliferative capacity until the FH2 stage, which may mark commitment to the hepatocyte lineage.

**Extended Data Fig. 2.**
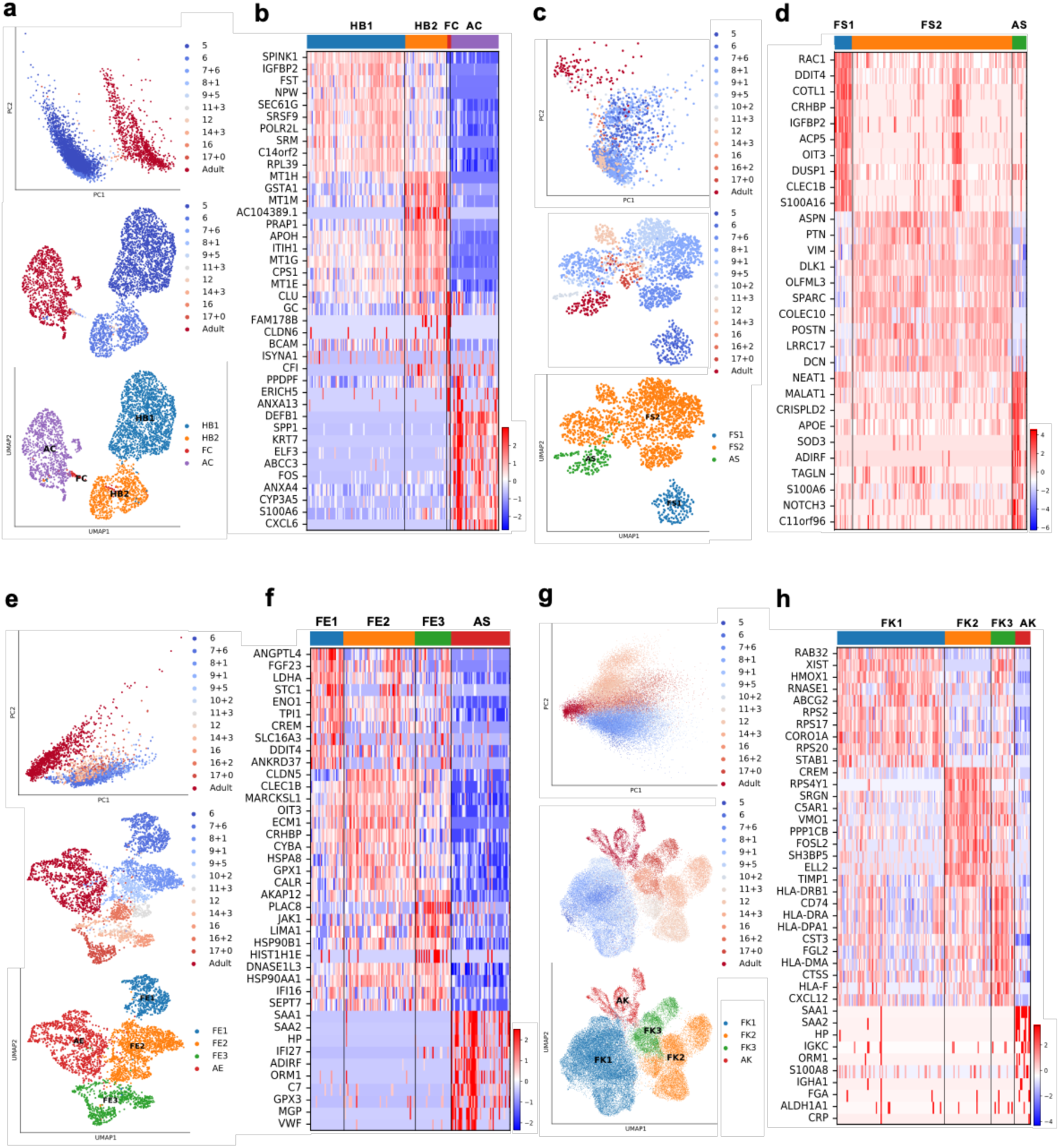
Mapping nonparenchymal cell identity during human liver development. **a,** PCA (top) and UMAP (middle) plots of primary human cholangiocyte sample timepoints and UMAP annotation of discrete cholangiocyte developmental stages (bottom); HB1, hepatoblast 1; HB2, hepatoblast 2; FC, fetal cholangiocyte; AC, adult cholangiocyte. **b,** Heatmap showing time-related DEGs of each stage of primary cholangiocyte development (Supplementary Table 4 for full list of DEGs). **c,** PCA (top) and UMAP (middle) plots of primary human hepatic stellate cell sample timepoints and UMAP annotation of discrete stellate cell developmental stages (bottom); FS1, fetal stellate cell 1; FS2, fetal stellate cell 2; AS, adult stellate cell. These three developmental stages coincide with the onset of haematopoietic function of the liver and birth. **d,** Heatmap showing time-related DEGs of each stage of primary Kupffer cell development (Supplementary Table 5 for full list of DEGs). **e,** PCA (top) and UMAP (middle) plots of primary human endothelial cell sample timepoints and UMAP annotation of discrete endothelial cell developmental stages (bottom); FE1, fetal endothelial cell 1; FE2, fetal endothelial cell 2; FE3, fetal endothelial cell 3; AE, adult endothelial cell. **f,** Heatmap showing time-related DEGs of each stage of primary Kupffer cell development (Supplementary Table 6 for full list of DEGs). Endothelial cells are closely associated with haematopoietic stem cell differentiation, with changes of function associated with haematopoietic and vascularization events. **g,** PCA (top) and UMAP (middle) plots of primary human Kupffer cell sample timepoints and UMAP annotation of discrete Kupffer cell developmental stages (bottom); FK1, fetal Kupffer cell 1; FK2, fetal Kupffer cell 2; FK3, fetal Kupffer cell 3; AK, adult Kupffer cell. **h,** Heatmap showing time-related DEGs specific to each stage of primary Kupffer cell development (Supplementary Table 7 for full list of DEGs).

**Extended Data Fig. 3.**
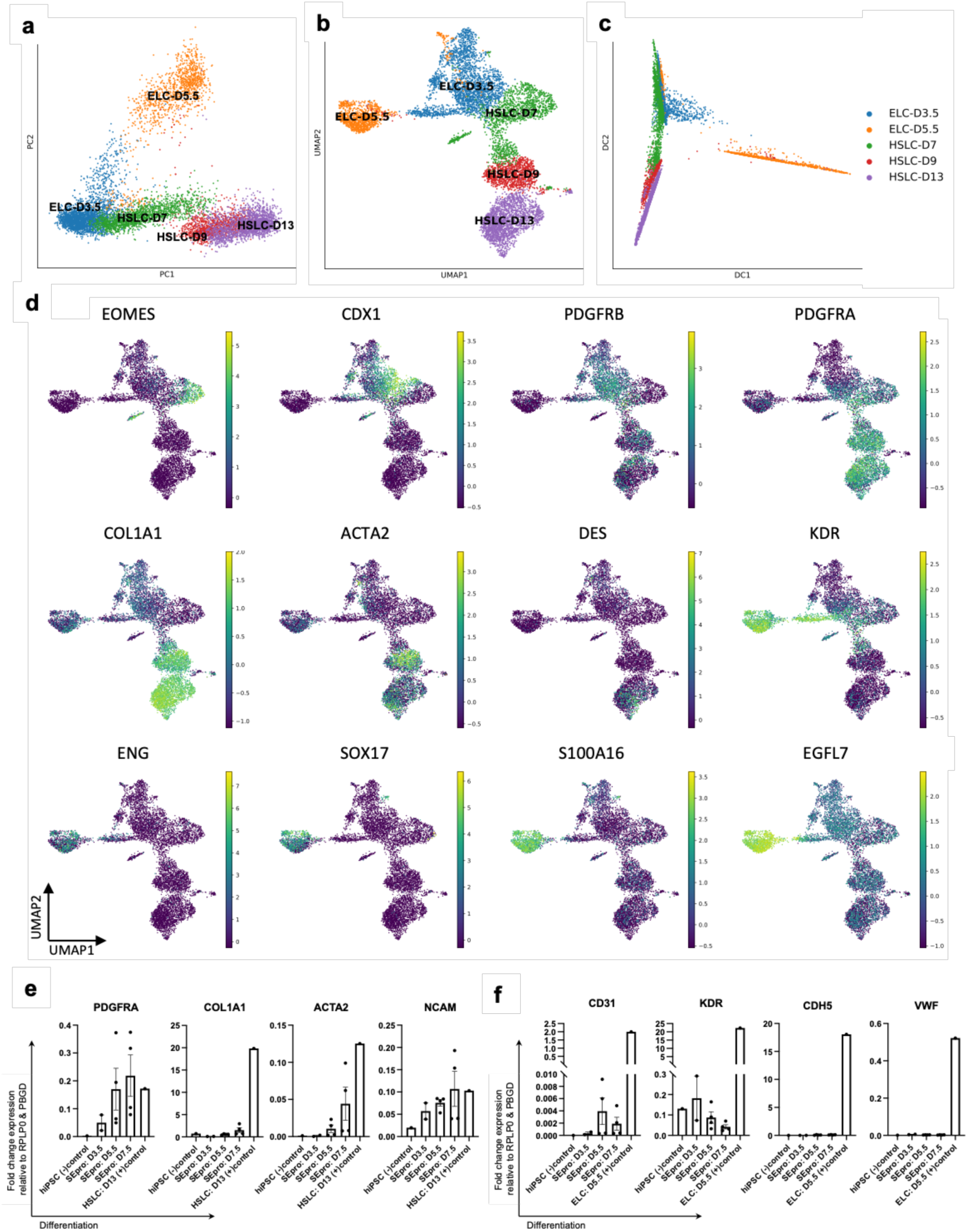
SEpro validation using an *in vitro* developmental model. **a,** PCA analysis of hiPSC-derived hepatic stellate cells (HSLC) and hiPSC-derived endothelial cells (ELC) confirming their common bipotent stellate-endothelial progenitor (SEpro) stage following mesoderm induction and before cell type specification. **b,** UMAP visualization using 30 neighbours and 500 PCs to confirm the strong lineage relationship of HSLCs and ELCs to the bipotent progenitor corresponding to the overlapping of ELC-D3.5 and HSLC-D7. **c,** Diffusion pseudotime confirming the specification of fetal-like HSLCs and ELCs from the common bipotent progenitor. **d,** UMAP visualizations showing the co-expression of key mesenchymal and endothelial markers such as CDX1, PDGFRB and KDR during the SEpro stage, followed by upregulation of lineage-specific markers and loss of co-expression. **e,** QPCR analyses of hiPSC differentiated first toward endothelial cells, then transitioned into culture conditions to specify hepatic stellate cells upon reaching the bipotential SEpro stage. These QPCR analyses show the acquisition of hepatic stellate cells markers and **f,** the loss of endothelial markers.

**Extended Data Fig. 4.**
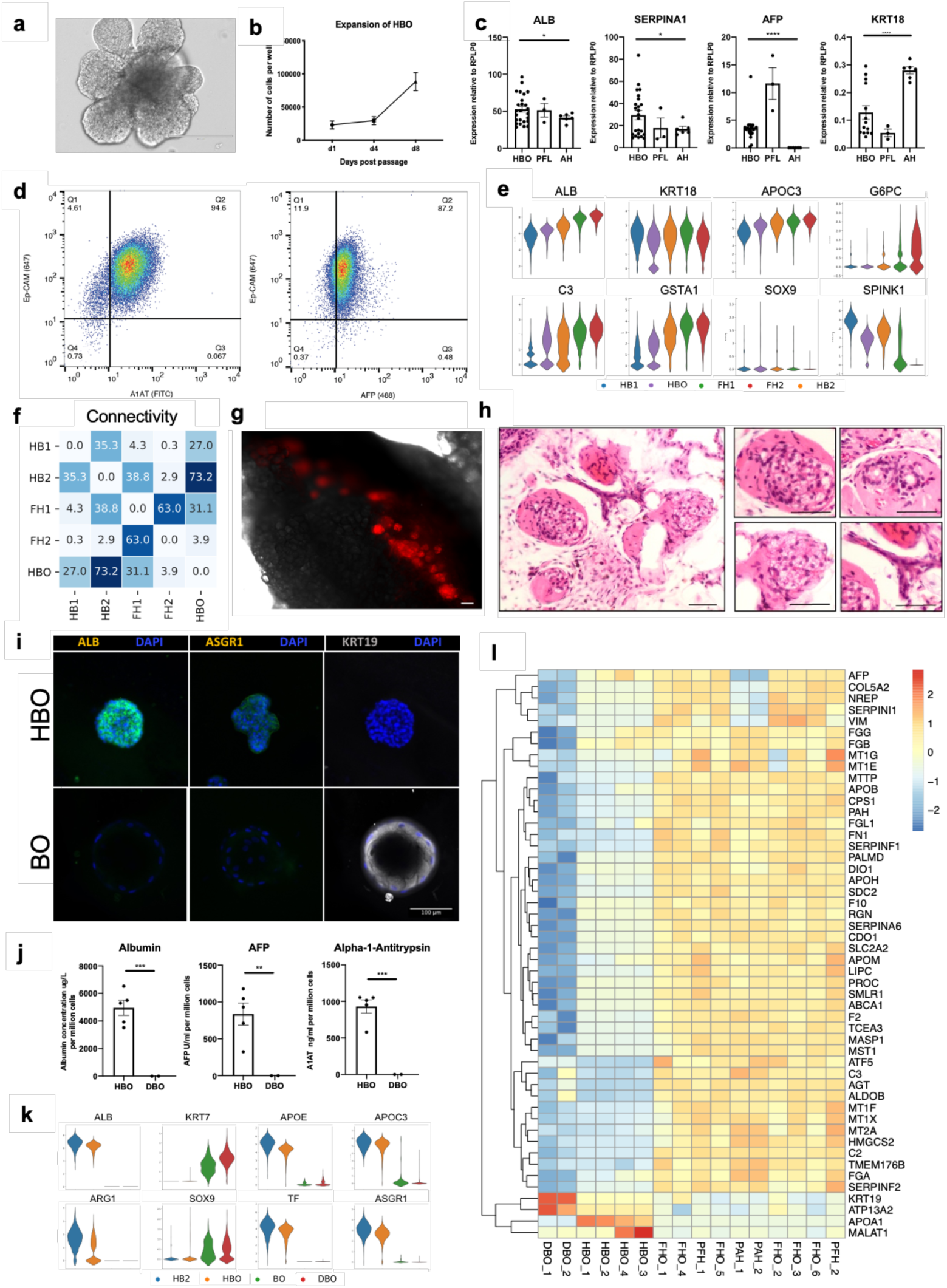
Characterisation of progenitor hepatoblast organoids and their unique developmental identity. **a,** Representative brightfield image of HBO. Scale bars = 200 um. **b,** Total number of cells per well averaged over three HBO lines at days 1, 4, and 8 post passage. **c,** Quantitative PCR analyses confirming the expression of key hepatoblast markers in HBO. Gene expression is shown as values relative to housekeeping gene RPLP-0. Each dot represents a biological replicate. HBO, n=25: (derived from n= primary 9 fetal livers at progressive passages from 3 to 15): ALB, SERPINA1, AFP, n=14: KRT18; primary fetal liver (PFL), n=3; primary adult hepatocytes (PAH), n=6. **d,** Fluorescence activated cell sorting (FACS) analysis on HBO (passage 11) using antibodies to EPCAM (647) and A1AT (488) (left), and EPCAM (647) and AFP (488) (right). **e,** ScRNA-seq violin plots comparing key functional markers of different stages of *in vivo* hepatocyte development to *in vitro* hepatoblast organoids, confirming their similarity to the HB2 stage. **f,** PAGA connectivity analysis confirming the resemblance of HBO and HB2. **g,** TdTomato-positive cell grafts (red) were identified in mouse fat (black, phase) upon explant of organoid tissues after 27 days of engraftment. **h,** Hematoxylin & Eosin staining of explanted grafts showing nodules (right, 4 images) with cells that morphologically resembled either densely packed hepatocytes (left) or biliary ductal-like structures (transverse section, right top; longitudinal section, right bottom). Scale bars = 100 um. **i,** Immunostaining of HBO and BO for albumin (ALB), asialoglycoprotein receptor 1 (ASGR1), and cytokeratin-19 (KRT19). Scale bar = 100 um. **j,** Concentration per litre of secreted proteins by HBO (n=5, each line derived from an independent fetal liver sample) and DBO (n=2, each line derived from an independent fetal liver sample) after 48 hours of freshly applied medium. Values are normalised to cell number (i.e. per million cells). **k,** ScRNA-seq violin plots highlighting the similarity in key hepatoblast functional gene expression between HB2 and HBO, and their dissimilarity to the cholangiocytic BO and DBO cultures. **l,** Heatmap of the top 50 absolute loadings in principal component one (32% variance) comparing DBO, FHO, HBO, PAH and PFH.

**Extended Data Fig. 5.**
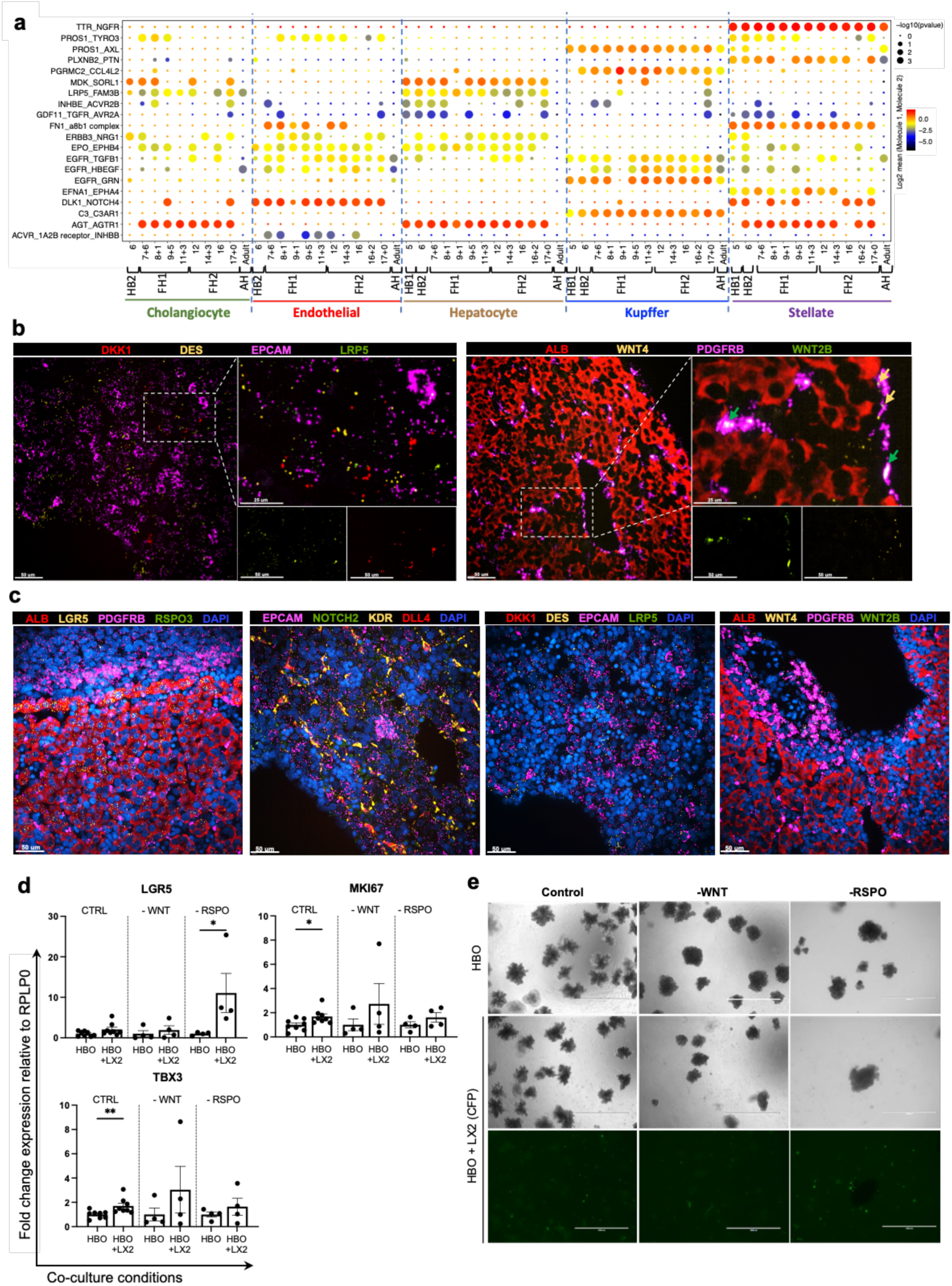
Identifying dynamic intercellular interactions during development that enable hepatocyte differentiation in the human liver. **a,** CellphoneDB analysis of hepatocytes receptors-ligand interactions with cholangiocytes, endothelial cells, hepatocytes, Kupffer cells and stellate cells across all developmental timepoints. These dynamic, temporal interactions with the nonparenchymal cells establish the hepatoblast/hepatocyte developmental extracellular environment. The Y-axis shows ligand-receptor/receptor-ligand interactions, with the hepatocyte protein listed first in the pair; x-axis shows developmental time of each interacting cell type; intensity shows log2 mean of interacting molecules; size of dot shows - log10(P) significance. **b,** RNAscope images showing DES+ hepatic stellate cells expressing DKK1 which interacts with the LRP5 receptor on EPCAM+ hepatoblasts (left panel). RNAscope images also reveal hepatoblasts expressing WNT4 and stellate cells expressing WNT2B, which is necessary for the proliferation and expansion of the early fetal liver hepatoblasts (right panel). **c,** RNAscope images including DAPI nuclear stain to visualise the co-expression of markers demonstrated by the RNAscope images in Fig. 4b and Extended Data Fig. 5b. Scale bars = 50 um.

**Extended Data Fig. 6.**
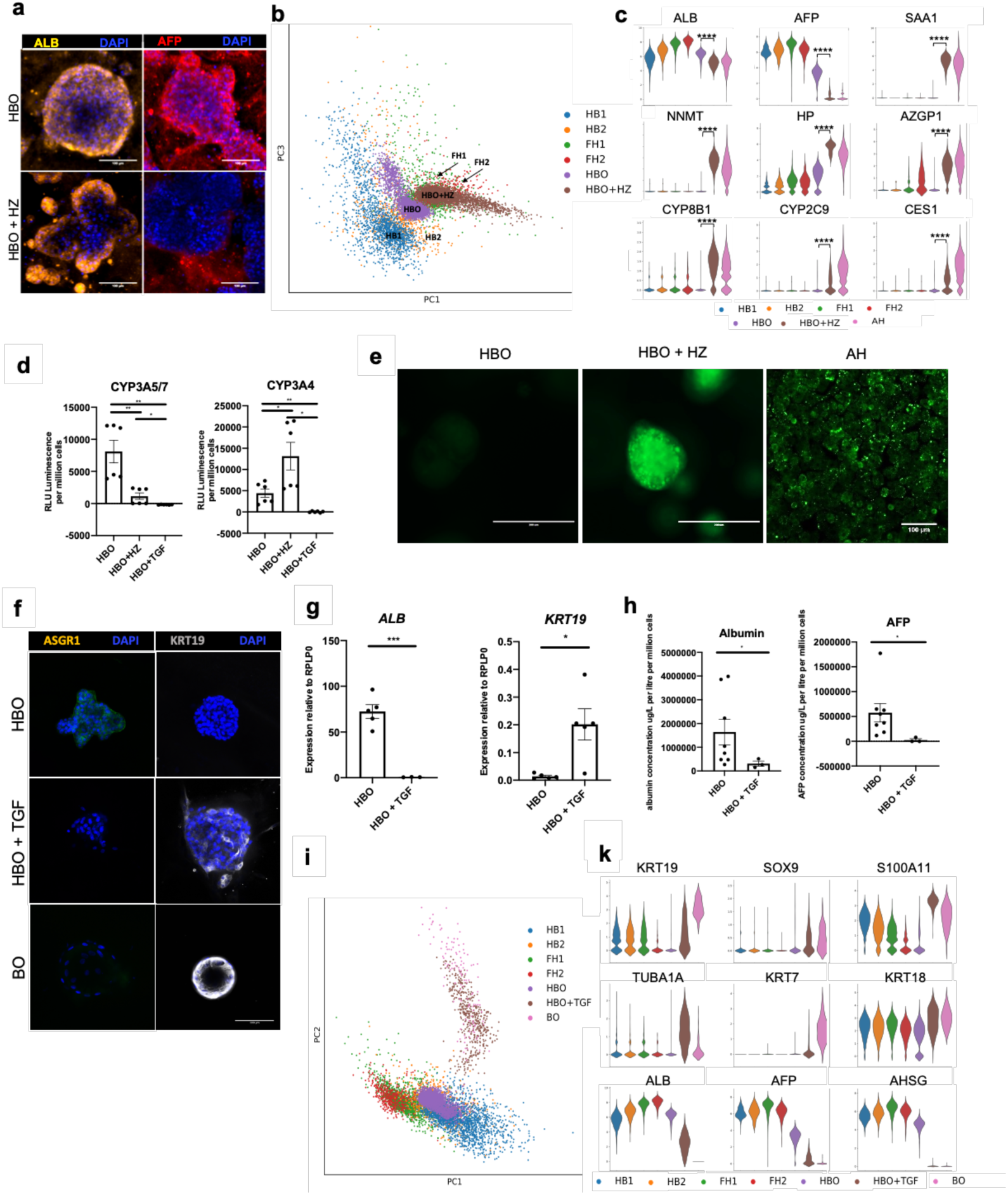
Characterisation of the differentiation capacity of hepatoblast organoids into both hepatocyte and cholangiocyte mature lineages. **a,** IF showing that HBO differentiated into hepatocytes maintain the expression of ALB while losing the foetal marker AFP. (HBO+HZ). **b,** PCA showing that HBO differentiation in vitro follows the developmental trajectory of foetal primary hepatocyte. **c,** Violin plots of key functional markers corresponding to the acquisition of an adult hepatocyte phenotype after HBO differentiation. *P<0.05, **P<0.01, ***P<0.001; two-tailed, unpaired t-tests. **d,** Cytochrome P450 3A5/7 and cytochrome P450 CYP3A4 activity in HBO, HBO+HZ, and HBO treated with TGFB (HBO+TGF). (lines derived from n=2 fetal livers, run in triplicate). **e,** BODIPY assay showing difference in lipid uptake in HBO compared to HBO + HZ and primary adult hepatocytes (PAH). Scale bars = 200 um. **f,** Immunocytochemistry of HBO, HBO+TGF, and BO stained for KRT19 and ASGR1. Scale bar = 100 um. **g,** QPCR analyses showing the expression of denoted genes in HBO and HBO+TGF; (n=5, each point represents an HBO line derived from a unique primary fetal liver). Gene expression is shown as relative value to housekeeping gene RPLP0. **h,** ELISA analyses showing the concentration of protein in media secreted by HBO (n=8, each line derived from an independent fetal liver sample) and HBO treated with TGFb (n=3) after 48 hours of freshly applied medium. Values are normalised to cell number (i.e. per million cells) with albumin as micrograms per litre, and alpha-fetoprotein as units per ml. **i,** PCA plot of scRNA-seq data comparing the *in vitro* differentiation of HBO toward cholangiocytes (HBO+TGF) to adult biliary organoids and *in vivo* differentiation of hepatoblasts. **j,** ScRNA-seq violin plots showing the loss of hepatocyte functional genes and the acquisition of a biliary transcriptome, thus demonstrating the similarity of HBO+TGF cholangiocytes to the positive biliary organoid control.

**Extended Data Fig. 7.**
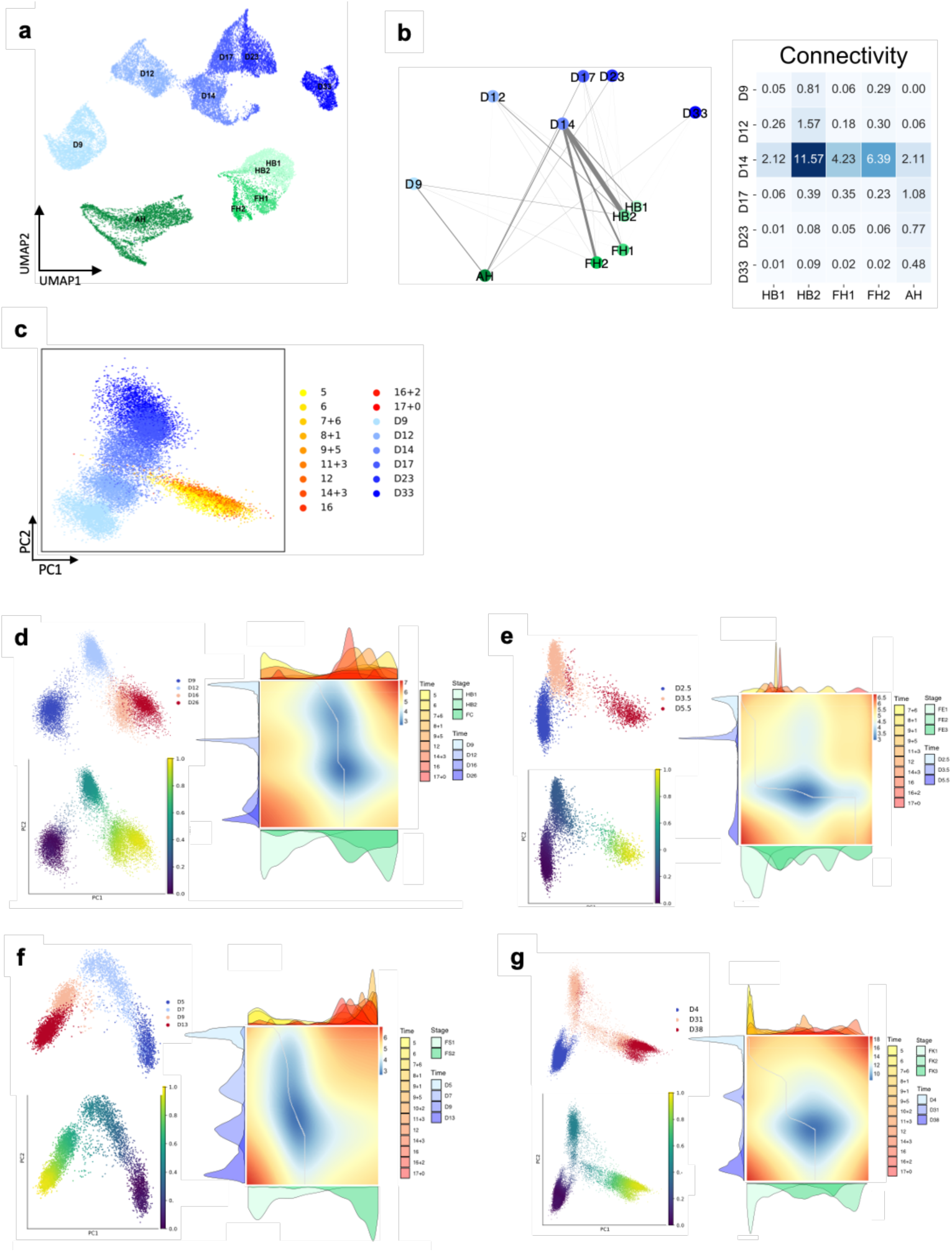
Comparison of hPSC-derived liver cell types to their primary counterparts during human liver development. **a,** UMAP visualisation showing primary hepatocyte development and hPSC-derived hepatocyte differentiation. **b,** PAGA connectivity plots and connectivity values comparing HLC timepoints to primary developmental timepoints/stages, confirming the similarity of the differentiation state of HLCs at day 14 (D14) to primary HB2. **c,** PCA plot of HLC diffusion map alignment to primary hepatoblast/hepatocyte development, validating the similarity of D14 HLC to HB2 shown in Fig. 5. **d,** PCA plot of cholangiocyte-like cells (CLCs) differentiation timepoints (top) and pseudotime (bottom) with 0.0 being the earliest pseudo-timepoint and 1.0 being the latest for all pseudotime analyses. Alignment of CLCs differentiation time course to primary cholangiocytes development (right), revealing the divergence of in vitro and in vivo differentiation at an early timepoint explaining the inability of CLCs to fully resemble primary adult cells. **e,** PCA plot of endothelial-like cells (ELC) differentiation timepoints (top) and pseudotime (bottom). Alignment of ELC differentiation time course to primary endothelial cell development (right) revealing the divergence of in vitro and in vivo differentiation at an early timepoint. **f,** PCA plot of hepatic stellate-like cells (HSLCs) differentiation timepoints (top) and pseudotime (bottom). Alignment of HSLCs differentiation time course to primary hepatic stellate cell development (right) revealing the divergence of in vitro and in vivo differentiation at an early timepoint. **g,** PCA plot of macrophage-like cells (MLC) differentiation timepoints (top) and pseudotime (bottom). Alignment of MLC differentiation time course to primary Kupffer cell development (right) revealing the divergence of in vitro and in vivo differentiation at an early timepoint.

**Extended Data Fig. 8.**
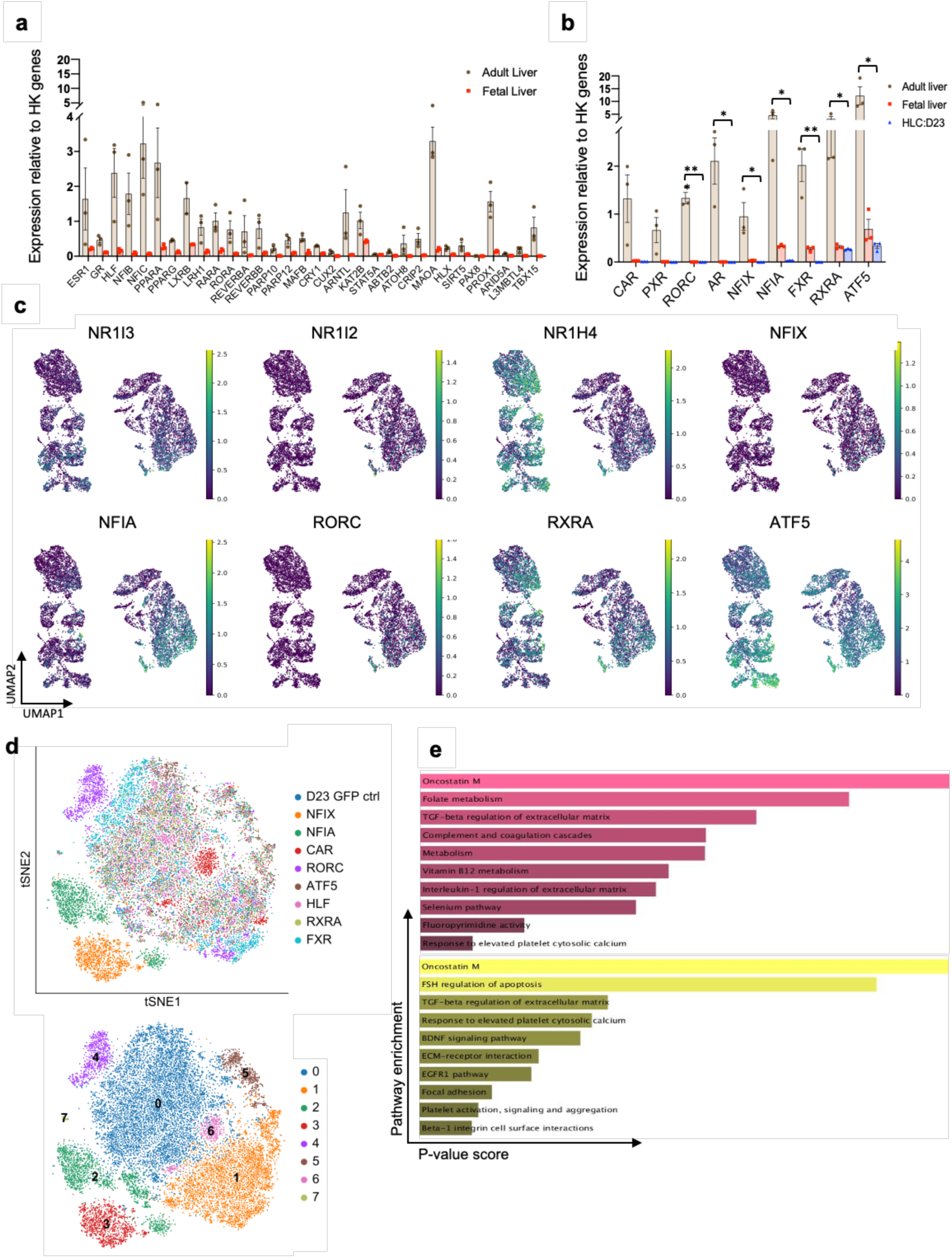
Improving *in vitro* functionality of hPSC-derived hepatocytes using the human liver developmental map. **a,** QPCR analyses showing the expression of transcription factors known to affect hepatocyte maturation in adult (n=3) and fetal (n=3, 6-10 PCW) human liver as well as **b,** factors identified from the scRNA-seq human liver development map as well as (Supplementary Table 15 for full lists); *P<0.05, **P<0.01, ***P<0.001; two-tailed, unpaired t-tests. **c,** UMAP visualization of transcription factors identified in Fig. 5 during hepatocyte development. Note that NFIX and NFIA (see Extended Data Fig. 1e) are expressed at low level in foetal stages and reach a maximum in adult hepatocytes. **d,** tSNE visualization of hPSCs hepatocytes transduced with selected transcription factors (TFs) or GFP indicting change in expressing profile. **e,** Pathway enrichment of top 150 DEGs comparing transduced HLCs to GFP control in **d** for NFIA (top) and NFIX (bottom) using the NCATS BioPlanet pathway database.

**Extended Data Fig. 9.**
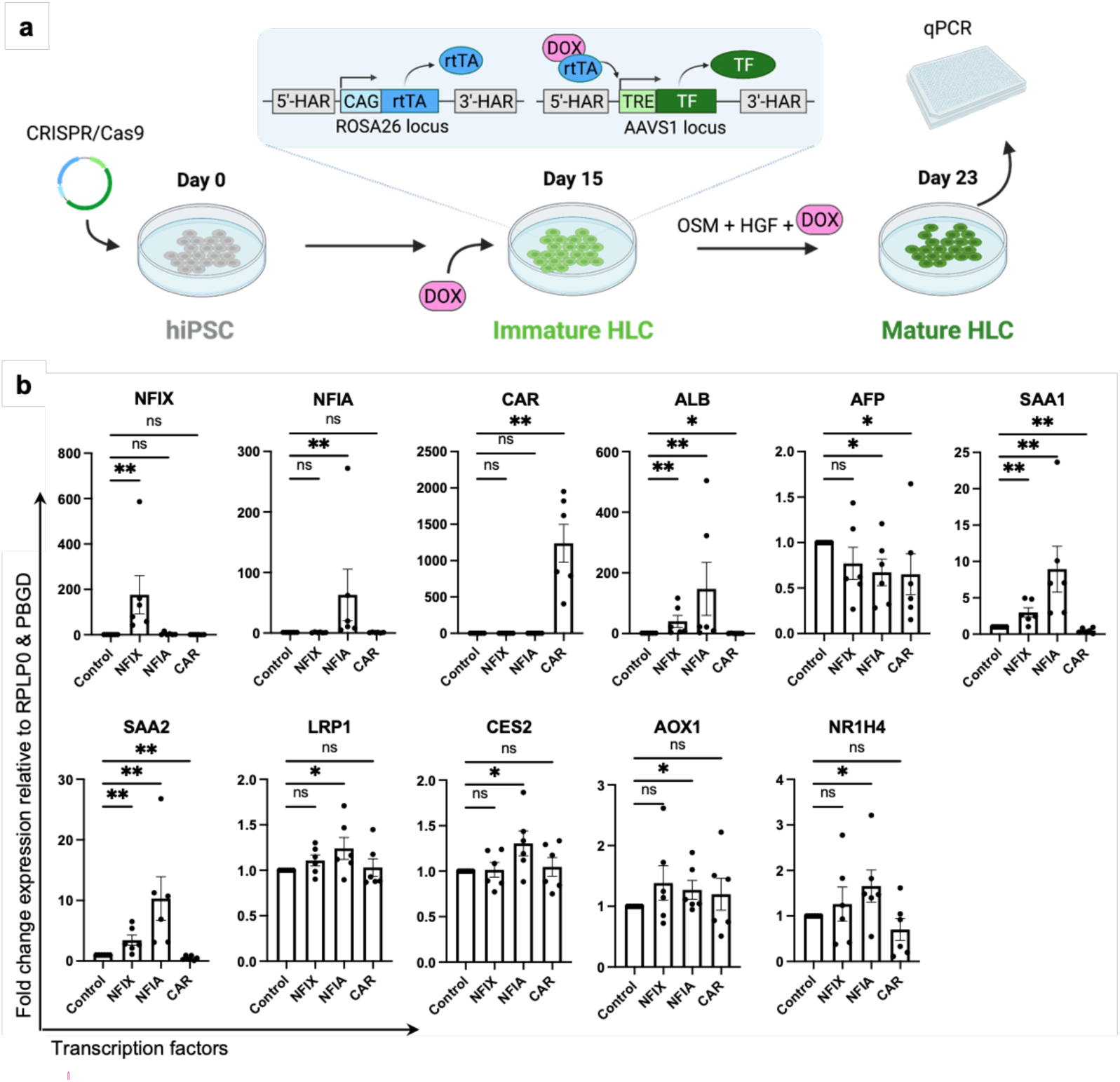
Validating the maturation of *in vitro* derived hepatocytes using an inducible expression culture system. **a,** Schematic of the experimental design for doxycycline (DOX) induction of key transcription factors during hepatocyte-like cell differentiation from hPSCs. HiPSC cells were edited to include the transcription factor in an inducible cassette, which was upregulated at day 15 of differentiation using DOX and continued to be expressed until day 23 when the HLCs were assays for maturation via qPCR. Created with BioRender.com. **b,** QPCR analysis of day 23 hepatocyte-like cells expressing NFIX, NFIA or CAR beginning at day 15 of differentiation using a doxycycline-inducible expression system. In addition to the increase in mature albumin expression and the decrease of fetal AFP expression, these results show the increase in a wide array of functional and metabolic hepatocyte markers, thus, indicating maturation due to temporally-relevant expression of key transcription factors. control: hepatocyte like cells grown in the absence of tetracycline. *P<0.05, **P<0.01, ***P<0.001; Mann-Whitney two-tailed, unpaired t-tests.

